# Genomic sequence characteristics and the empiric accuracy of short-read sequencing

**DOI:** 10.1101/2021.04.08.438862

**Authors:** Maximillian Marin, Roger Vargas, Michael Harris, Brendan Jeffrey, L. Elaine Epperson, David Durbin, Michael Strong, Max Salfinger, Zamin Iqbal, Irada Akhundova, Sergo Vashakidze, Valeriu Crudu, Alex Rosenthal, Maha Reda Farhat

## Abstract

**Background:** Short-read whole genome sequencing (WGS) is a vital tool for clinical applications and basic research. Genetic divergence from the reference genome, repetitive sequences, and sequencing bias, reduce the performance of variant calling using short-read alignment, but the loss in recall and specificity has not been adequately characterized. For the clonal pathogen Mycobacterium tuberculosis (Mtb), researchers frequently exclude 10.7% of the genome believed to be repetitive and prone to erroneous variant calls. To benchmark short-read variant calling, we used 36 diverse clinical Mtb isolates dually sequenced with Illumina short-reads and PacBio long-reads. We systematically study the short-read variant calling accuracy and the influence of sequence uniqueness, reference bias, and GC content. å

**Results:** Reference based Illumina variant calling had a recall ≥89.0% and precision ≥98.5% across parameters evaluated. The best balance between precision and recall was achieved by tuning the mapping quality (MQ) threshold, i.e. confidence of the read mapping (recall 85.8%, precision 99.1% at MQ ≥ 40). Masking repetitive sequence content is an alternative conservative approach to variant calling that maintains high precision (recall 70.2%, precision 99.6% at MQ≥40). Of the genomic positions typically excluded for Mtb, 68% are accurately called using Illumina WGS including 52 of the 168 PE/PPE genes (34.5%). We present a refined list of low confidence regions and examine the largest sources of variant calling error.

**Conclusions:** Our improved approach to variant calling has broad implications for the use of WGS in the study of Mtb biology, inference of transmission in public health surveillance systems, and more generally for WGS applications in other organisms.

## Background

Illumina short-read whole genome sequencing (WGS) followed by alignment to a reference genome is widely used to identify genetic variants. Illumina sequencing and alignment can confidently detect single nucleotide substitutions (SNSs) and small insertions or deletions (INDELs) but is limited in several ways by its short ~100 bp target read lengths. First, short repetitive or homologous query sequences are challenging to uniquely align to the genomic reference^1,2^. Second, genomic DNA extraction and sequencing library preparation of short-reads may be more error or bias prone^3–7^. For example, regions with high GC content and/or low sequence complexity may be particularly prone to PCR-dropout and reduced sequencing coverage^7–9^. Third, the use of a single reference genome introduces bias, especially when the genome being analyzed differs substantially from the reference sequence^10,11^. As the sequenced genome diverges from the reference genome, short-read alignment becomes increasingly inaccurate and regions absent from the reference genome are missed or poorly reconstructed.

In contrast, long-read sequencing can generate high confidence complete genome assemblies, which can also be used to benchmark Illumina WGS. For example, long-reads generated by PacBio sequencing (with lengths on the order of ~10 kb) are ideal for assembling complete bacterial genomes and identifying variants in repetitive regions^12^. Although individual PacBio reads have a considerably higher per base error rate (10-15%) than Illumina, the randomly distributed nature of the errors allows for high coverage sequencing runs to converge to a high accuracy consensus^13^. More recently, circular consensus sequencing has further improved PacBio long-read per base accuracy to levels on par with Illumina^14^. Alternatively, hybrid strategies that combine less accurate long-reads and short Illumina reads can offer both high base-level accuracy and continuity of the final assembly^12,15^.

*Mycobacterium tuberculosis* (Mtb) is a globally prevalent pathogenic bacterium with a ~4.4 Mbp genome known for high GC content, large repetitive regions, and an overall low mutation rate. Owing to the clonality and stability of the Mtb genome, this organism is particularly well suited for systematically identifying the sources of error that arise when short-read data is used for variant detection. Approximately 10% of the Mtb reference genome (H37Rv) is regularly excluded from genomic analysis because it is purported to be more error prone and enriched for repetitive sequence content^16^. This 10% of the Mtb genome, hitherto regions of putative low confidence (PLC), span the following genes/families: 1) PE/PPE genes (N=168), 2) mobile genetic elements (MGEs) (N=147), and 3) 69 additional genes with identified homology elsewhere in the genome^17^. Despite their systematic exclusion from most Mtb genomic analyses^17–19^, PLC regions are yet to be evaluated systematically for short-read variant calling accuracy. Here, we use long-read sequencing data from 36 phylogenetically diverse Mtb isolates to benchmark short-read variant detection accuracy and study genome characteristics that associate with erroneous variant calls.

## Results

### High confidence Mtb assemblies with hybrid short- and long-read sequencing

For this study, PacBio long-read and Illumina sequencing was performed for 31 clinical Mtb isolates. The resultant data was combined with publicly available paired PacBio and Illumina genome sequencing of 18 Mtb isolates from two previously published studies^20,21^. From these datasets, a total of 38 clinical isolates were selected for having a) paired end Illumina WGS with median sequencing depth ≥ 40× relative to the Mtb reference genome, and b) no evidence of mixed infections or sample swaps (**Additional File 2**).

Across these 38 isolates, the mean sequencing depth relative to the H37Rv reference genome was 84× (IQR: 67× − 107×) for Illumina and 286× (IQR: 180× − 367×) for PacBio. We performed *de novo* genome assembly and iteratively polished each assembly with the PacBio and Illumina reads generating a complete circular assembly for 36/38 isolates (**Methods**). For uniformity in assembly completeness, we excluded the 2 non-circular assemblies from downstream analysis.

We assessed the accuracy of the *de novo* PacBio assemblies by examining the profile of errors corrected during the Illumina polishing step (**Supp. Figure 1, Additional File 3**). Across all 36 assemblies, erroneous 1-bp insertions and deletions (INDELs) made up 97.9% of all corrections made by Illumina polishing with Pilon^22^. The median number of erroneous insertions and deletions per assembly was 5 (IQR: 2 - 88) and 15 (IQR: 4 - 37) respectively. Very few of the errors corrected during Illumina polishing were single nucleotide changes; median of 0 (IQR: 0 - 2) across all polished 36 genome assemblies. Overall, the number of changes made during Illumina polishing of the *de novo* PacBio assembly was negatively correlated to PacBio sequencing depth (Spearman’s R = −0.458, p < 4.9e-3) (**Supp. Figure 1C**).

The 36 assemblies spanned the Mtb global phylogeny and had a high degree of conservation in genome structure and content relative to the H37Rv reference genome (**Figure 1, Supp. Figure 2**): Average Nucleotide Identity (ANI) to H37Rv (99.84% to 99.95%), genome size (4.38-4.44 Mb), GC content (65.59 - 65.64%), and predicted gene count (4017 - 4096 ORFs) (**Additional File 2**).

**Figure 1.**
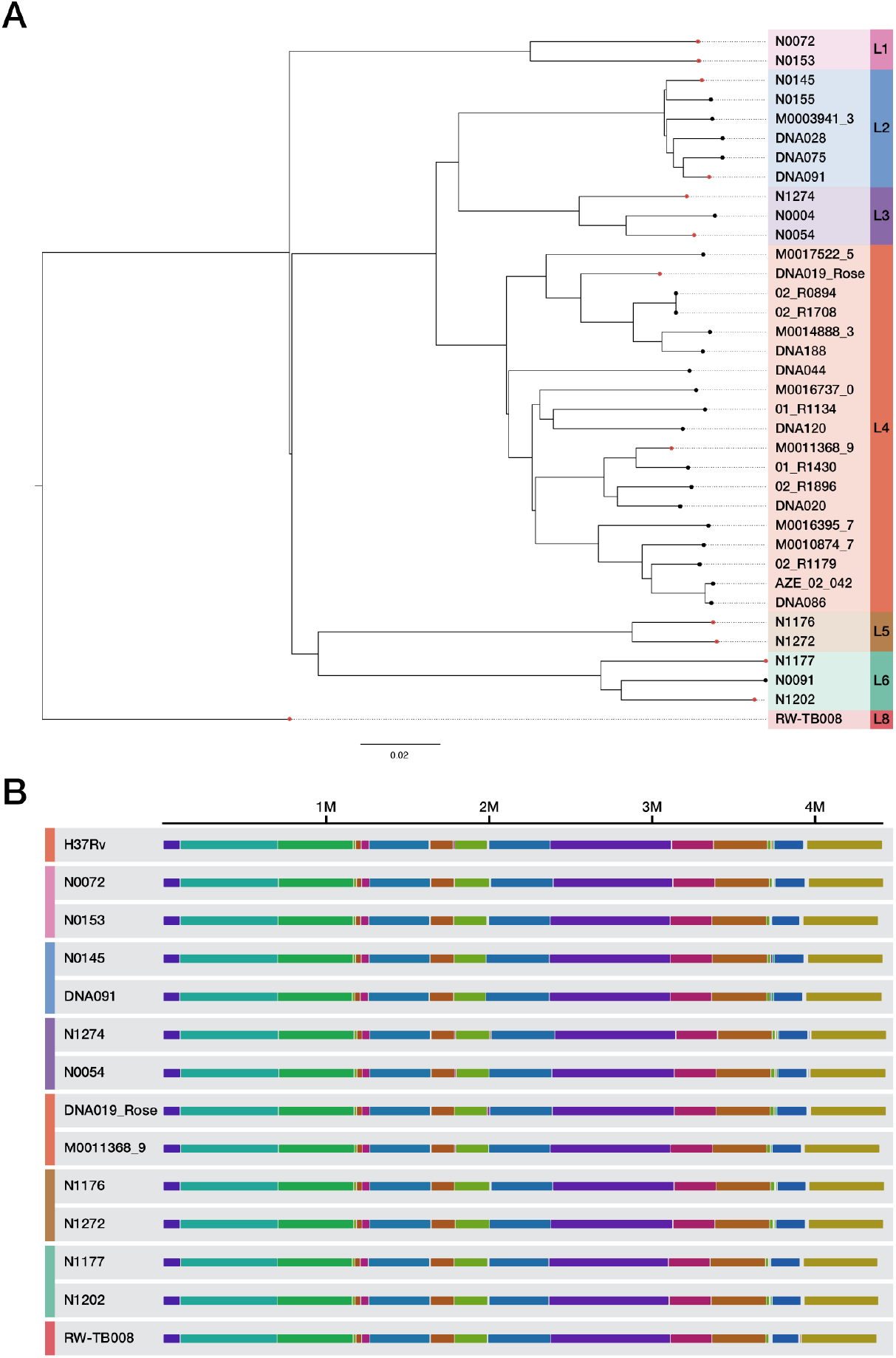
Overview of 36 clinical Mtb isolates with completed genome assemblies. **a)** Maximum likelihood Phylogeny of *M. tuberculosis* isolates with PacBio complete genome assemblies. The sequences of all 36 complete *M. tb* genomes were aligned to the H37rv reference genome using minimap2, and a maximum likelihood phylogeny was inferred using a concatenated SNS alignment (15,673 total positions). **b**) Representative isolates from each lineage sampled from the whole genome sequence alignment between the H37Rv reference genome and all completed circular Mtb genome assemblies, The complete alignment is visualized in Supplemental Figure 2. The whole genome multiple sequence alignment was performed using the *progressiveMauve*^44^ algorithm. Each contiguously colored region is a locally collinear block (LCB), a region without rearrangement of homologous backbone sequence.

In accordance with the small variant benchmarking guidelines of Global Alliance for Genomics & Health^23^(GA4GH), we excluded a small subset of regions with ambiguous ground truths on a per isolate basis (**Methods**). These ambiguous regions fell into 2 categories: a) variable copy number relative to the H37Rv reference genome or b) difficult to align regions due to a high level of sequence divergence relative to the reference genome. We excluded these regions from our performance evaluation in this paper due to their difficulty of interpretation **(Additional File 4)**. The percentage of the genome identified as ambiguous was consistently lower than 1% (median: 0.41%, IQR: 0.28% - 0.49%) across all assemblies. We observed that for the regions that were frequently ambiguously (Ambiguous in > 25% of isolates, **Additional File 5)**, 96.8% of bases were from regions which overlapped with recognized PLC regions.

### Empirical base-level performance of Illumina

To measure the consistency and accuracy of Illumina genotyping across the Mtb genome, we defined the Empirical Base-level Recall metric (EBR) for each position of the H37Rv reference genome (4.4 Mb, **Additional File 6**). EBR was calculated as the proportion of isolates for which Illumina variant calling made a *confident* variant call that agreed with the ground truth, hence a site with a perfect (1.0) EBR score requires Illumina read data to pass the default quality criteria (**Methods**), and then agree with the PacBio defined ground truth for 100% of the isolates (Examples in **Figure 2**). EBR was significantly lower within PLC regions (mean EBR = 0.905, N = 469,501 bp) than the rest of the genome (mean EBR = 0. 998, N = 3,942,031 bp, Mann-Whitney U Test, P < 2.225e-308) (**Figure 3A, Table S1**). But EBR was not consistently low across PLC regions, with 67% of PLC base positions having EBR ≥ 0.97. EBR averaged by gene (gene-level EBR) also showed heterogeneity across PLC regions with 62.6%, 61.3% and 82.6% respectively of the MGEs, PE/PPE, and previously classified repetitive genes having gene-level EBR ≥ 0.97 (**Figure 3B, Supp. Figure 3, Tables S2-S3, Additional File 7).** All other, non-PLC, functional gene categories had a median gene<u>-</u>level EBR =1, among these only 14 non-PLC genes had a gene-level EBR < 0.97.

**Figure 2.**
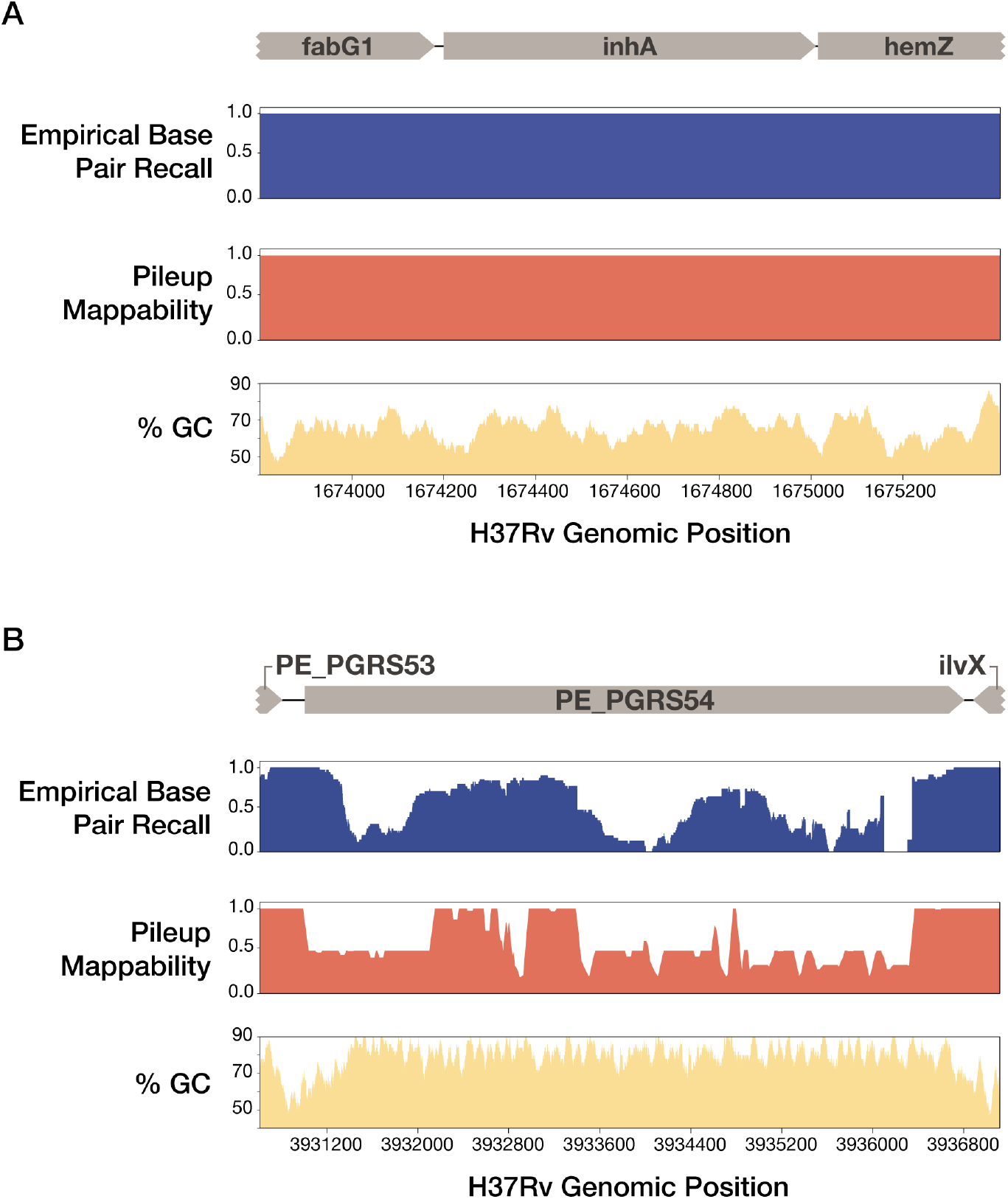
EBR, Pileup Mappability, and GC content across two example regions of the H37Rv genome. Empirical Base Pair Recall (EBR), Pileup Mappability (K=50 bp, e = 4 mismatches) and GC% (100 bp window) values are plotted across all base pair positions of two regions of interest. **a)** InhA, an antibiotic resistance gene, shows perfect EBR across the entire gene body. **b)** In contrast, PE_PGRS54, a known highly repetitive gene with high GC content, has extremely low EBR across the entire gene body. Browser tracks of EBR and Pileup Mappability in BEDGRAPH format are made available as Additional Files 17 and 18.

**Figure 3.**
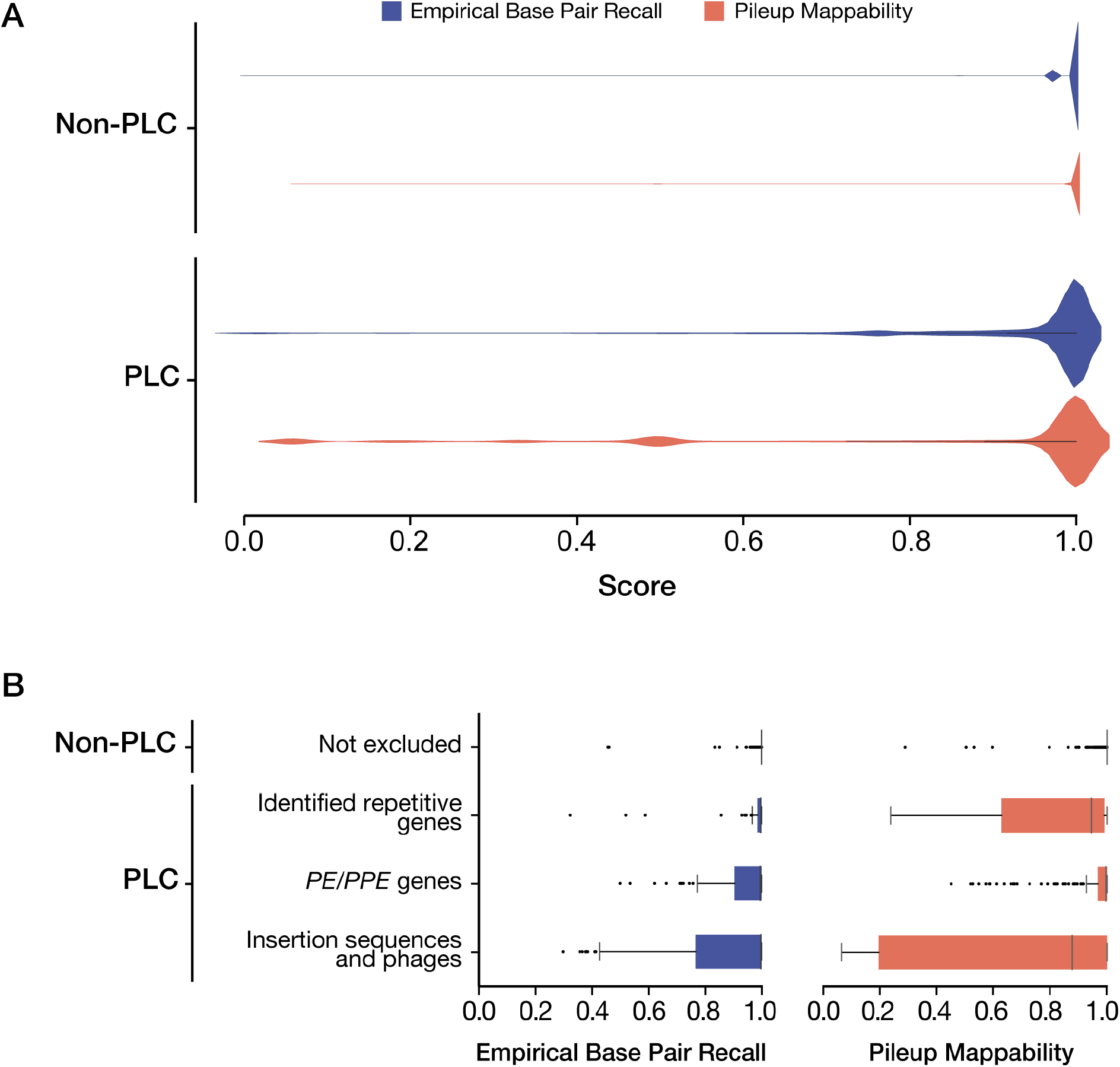
The Distribution of EBR and Pileup Mappability scores in PLC and non-PLC regions. **a)** The distribution of Empirical Base Pair Recall (EBR) and Pileup Mappability (P-Map, K=50,E=4) scores of PLC and non-PLC regions. Excluded regions harbor significantly more low EBR base pair positions when compared to the included genes, but 68% of routinely excluded positions still have ≥ 97% EBR. The Pileup mappability with K=50 bp is lower in PLC regions (mean = 0.86) than non-PLC regions (mean = .997). **b)** The Distribution of gene-level mean EBR and P-Map (K=50,E=4) between PLC and non-PLC regions. We compared the mean EBR and Pileup Mappability across all genes within PLC and non-PLC regions. The pe and ppe gene families (PE/PPEs) and mobile genetic elements (MGE), which make up 82% of PLC genes, demonstrated significantly lower mean EBR and Pileup Mappability than other non-PLC genes.

### Characteristics of regions with low empirical performance

Across all 36 isolates evaluated, we observed 1,825,385 sites where Illumina failed to confidently agree with the inferred ground truth. These low recall sites were spread across 267,471 unique positions of the H37Rv reference genome with EBR < 1. We explored the underlying factors associated with low recall at these positions using the associated filter and quality tags provided by the variant caller, Pilon (**Methods, Table S4**). Across the 1,829,181 low recall sites, the distribution of outcomes included: a) 62.78% low coverage (LowCov), b) 30.74% falsely called as deleted (Del) with or without low coverage or other tags, c) 6.24% were missed deletions tagged as PASS, d) 0.03% (669 sites) were false base calls (reference or alternate) tagged as PASS, e) 0.25% remaining positions were labeled as ambiguous (Amb) due to evidence for two or more alleles at a frequency ≥ 25%.

Among all low recall sites annotated as with a Low Coverage tag: (a) 45.8% were due to insufficient total coverage of aligned reads (sequencing bias or extreme sequence divergence, total Depth < 5), (b) 27.6% lacked uniquely aligning reads (repetitive sequence content, mapping quality = 0), and (c) 26.6% were due to low confidence paired-end alignments that did not pass Pilon’s heuristics (likely structural variation causing improper paired-alignment orientation).

### Repetitive sequence content

We identified repetitive regions in H37Rv and evaluated their relationship with low EBR using the pileup mappability metric (**Methods**). Pileup mappability scores range from 0 to 1, where 1 represents a genomic position where all overlapping sequence K-mers are unique in the genome of interest within a similarity threshold of E mismatches. We calculated pileup mappability conservatively with a K-mer size of 50 base pairs and up to 4 mismatches (P-Map-K50E4, **Additional File 6)**. P-Map-K50E4 is lower in PLC regions (mean = 0.856) than non-PLC regions (mean = .997), (Mann-Whitney U Test, P < 0.001) (**Figure 3A**). Yet, 69.7% of positions in PLC regions had P-Map-K50E4 scores of 1, indicating uniquely alignable sequence content even with sequence lengths as short as 50 bp (**Table S5**). At the gene-level, PE/PPEs and MGEs had lower P-Map-K50E4 than the rest of the genome (Wilcoxon, P < 2e-308) (**Figure 3B, Table S6, Additional File 7**) but 34.5%, and 32.7% of these genes respectively had perfect (1.0) P-Map-K50E4 across the entire gene body. Previously identified repetitive genes (N = 69) had a gene-level P-Map-K50 below 1 which is expected given that this was their defining feature^24^, but for the majority (51 of 69), median mappability was greater than 0.99, indicating that a high proportion of their sequence content was actually unique. Non-PLC functional categories had a median gene level P-Map-K50E4 = 1.0 (**Supp. Figure 3, Table S7**). Genome-wide P-Map-K50E4 and EBR scores were moderately correlated (Spearman’s *ρ*= 0.47, P < 2e-308). Thirty percent of all genome positions with EBR < 1.0 also had a P-Map-K50E4 score below 1.0.

### Sequencing bias in high GC-content regions

Across several sequencing platforms, high-GC content associates with low sequencing depth due to low sequence complexity, PCR biases in the library preparation and sequencing chemistry^3–6^. We assessed the sequencing bias of Illumina and PacBio across each individual genome assembly using the relative depth metric^4^(the depth per site divided by average depth across the entire assembly) to control for varying depth between isolates. On average with Illumina, 1.2% of the genome had low relative depth (< 0.25), while for PacBio sequencing the average proportion of the genome with low relative depth was 0.0058% (Mann-Whitney U Test, P < 0.001). Both sequencing technologies demonstrated coverage bias against high-GC regions, with more extreme bias for Illumina than PacBio (**Figure 4, Additional File 8**). Across all base pair positions with local GC% ≥ 80%, using a window size of 100 bp, the mean relative depth was 0.79 for PacBio and 0.35 for Illumina. Genome-wide, EBR was significantly negatively correlated with GC content (Spearman’s *ρ*= − 0.12, P < 2e-308), but this correlation was weaker than that observed with sequence uniqueness (P-Map-K50E4, as above Spearman’s *ρ*=0.47).

**Figure 4.**
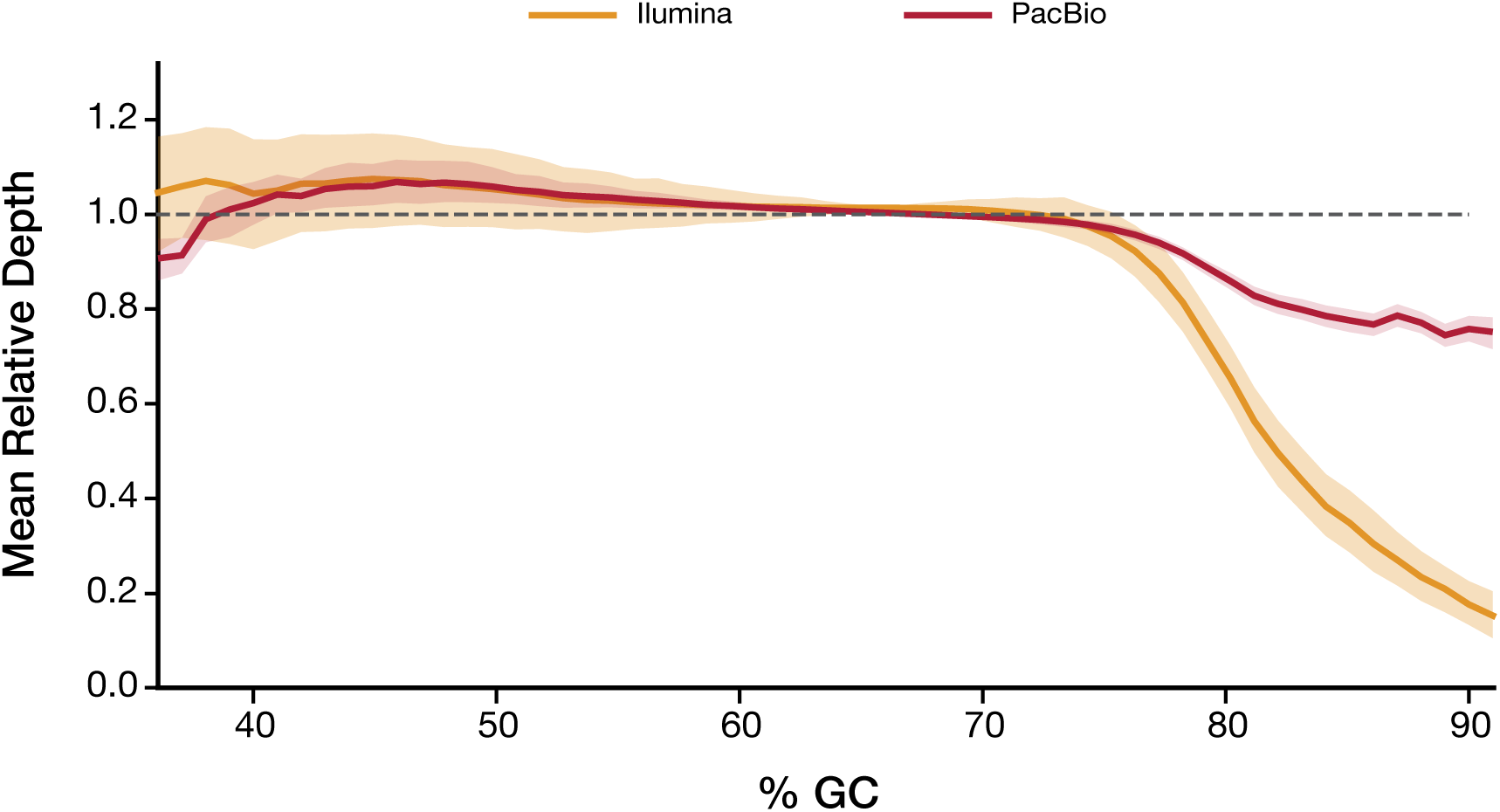
Relative sequencing depth as a function of local GC content across all 36 complete isolates. We evaluated the relative depth of our Illumina and PacBio sequencing data as a function of GC content (100 bp window size) across all positions of each isolate’s complete genome assembly. The relative depth was averaged across all positions with the same GC% across each genome assembly. The standard error of the mean of the relative depth across all 36 isolates is shaded for each sequencing technology. At high (>70%) GC contents, Illumina starts to show lower relative depth compared to PacBio sequencing.

### False positive SNS variant calls

Next, we focused specifically on regions with high numbers of false positive SNSs identified through comparison with the ground-truth variant calls. We examined the distribution of false positive SNS calls across the H37Rv reference genome using a realistic intermediate variant filtering threshold of mean mapping quality at the variant site (MQ ≥ 30, **Figure 5, Additional File 9**). The top 30 regions ranked by the number of false positives (23 genes and 7 intergenic regions) contained 89.4% (490/548) of the total false positive calls and spanned 65 kb, 1.5% of the H37Rv genome. Of these 30 false positive hotspot regions, 29 were either a PLC gene or an intergenic region adjacent to a PLC gene: 17 PE/PPE genes, 3 MGEs, 2 were previously identified repetitive genes^24^, and 7 PLC-adjacent intergenic regions. Across all false positives, the PE-PGRS and PPE-MPTR sub-families of the PE/PPE genes were responsible for a large proportion (45.4%) of total false positive variant calls. Of all the 556 false positives SNSs evaluated (MQ ≥ 30), only 14 were detected across 4 non-PLC genes: Rv3785 (9 FPs), Rv2823c (1 FP), plsB2 (2 FPs), Rv1435c (2 FPs).

**Figure 5.**
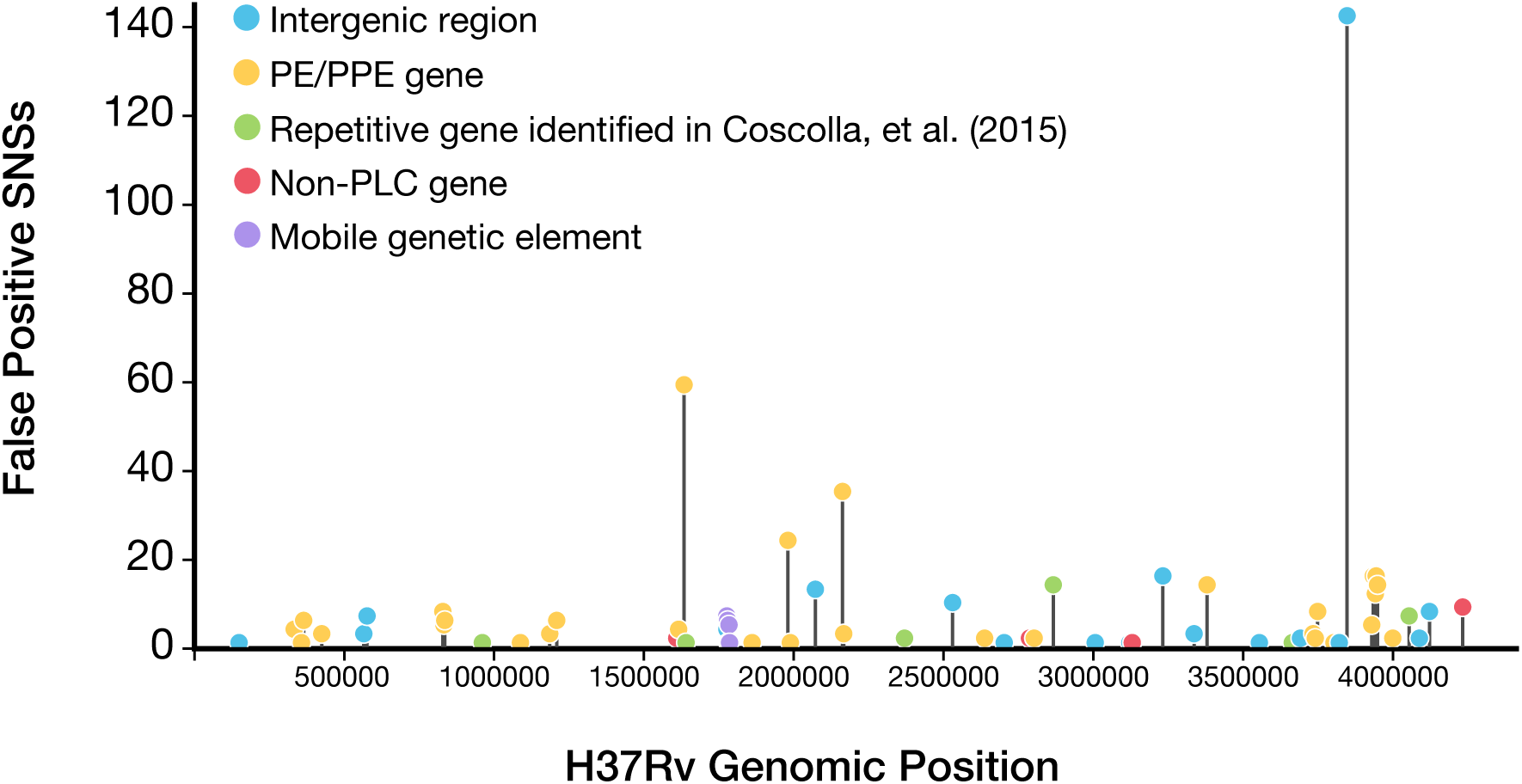
The distribution of potential false positive SNS calls across all genomic regions of the H37Rv genome. The frequency of false positive SNS calls detected (MQ ≥ 30) across all 36 isolates evaluated was plotted for all regions of the H37Rv genome (gene or intergenic regions). The top 30 regions ranked by the number of total false positives contained 89.4% (490/548) of the total false positive SNSs and spanned only 65 kb of the H37Rv genome. Full results for all annotated genomic regions (gene or intergenic) can be found in Additional File 9.

### Masking to balance precision and recall

A common approach for reducing Mtb false positive variant calls is to mask/exclude all PLC regions from variant calling. Here we investigated two variations on this that utilize directly reference sequence uniqueness and variant quality metrics. We compared: (1) masking of regions with non-unique sequence, defined as positions with P-Map-K50E4 < 1, (2) No *a priori* masking of any regions, and (3) masking of all PLC genes (the current standard practice). We then filtered potential variant calls by whether the variant passed all internal heuristics of the Pilon^22^-based variant calling pipeline (**Methods**) and studied the effect of varying the mean mapping quality (MQ) filtering threshold from 1 to 60 (**Figure 6**). We computed the F1-score, precision and recall of detection of SNSs and small indels (<=15bp) for each masking schema and MQ threshold across all 36 clinical isolates (**Methods, Additional File 10**).

**Figure 6.**
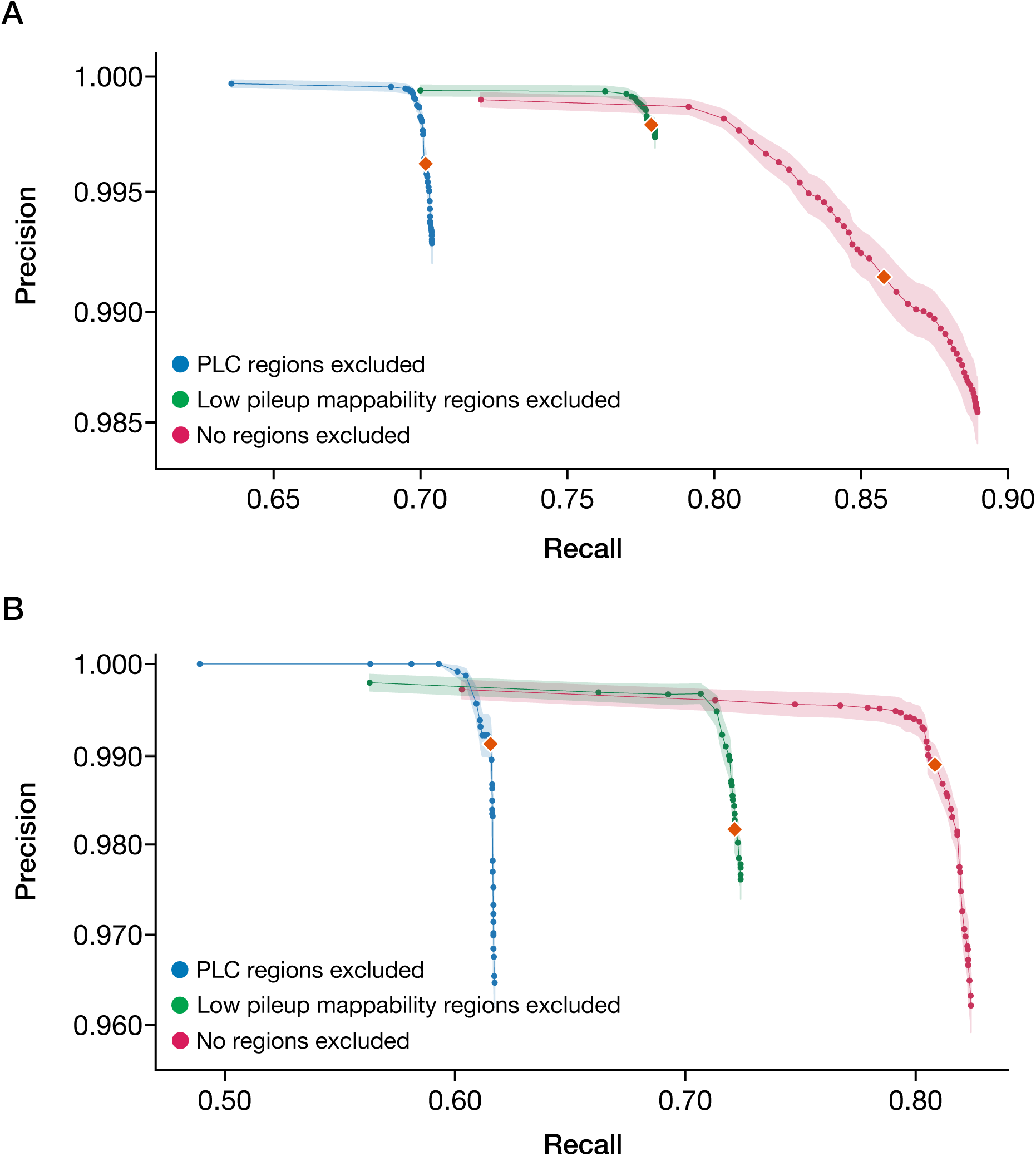
Mean SNV and INDEL variant calling performance across different masking approaches. **a)** SNS variant calling performance was evaluated across the following three schemas: (1) masking of regions with non-unique sequence, as defined as positions with P-Map-K50E4 < 1, (2) No *a priori* masking of any regions, and compared to (3) masking of all PLC genes (the current standard practice). (**b)** short INDEL (1-15 bp) variant calling performance was evaluated across the same schemas. The orange diamonds represent the mean precision and recall using a MQ threshold of 40 for both (a) and (b). Shaded regions represent the SEM of precision across all 36 isolates evaluated. For all masking approaches evaluated, the MQ thresholds evaluated ranged from 1-60. Complete benchmarking results can be found for each individual isolate in Additional File 10.

For SNSs, mean recall ranged from 63.6% to 89.0%, and precision ranged from 98.5% to 99.97% across the three schemas (**Figure 6A**). At a threshold of MQ ≥ 40, we observed the following mean SNS performances: 1) Masking non-unique regions, F1 = 0.87 (Precision = 99.8%, Recall = 77.9%), 2) no masking of the genome, F1 = 0.92 (Precision = 99.1%, Recall = 85.8%), 3) Masking PLC genes, F1 = 0.82 (Precision = 99.6%, Recall = 70.2%). Based on F1 score, no masking of the genome had the highest overall performance, but masking non-unique regions had the highest precision. Decreasing the MQ threshold to an optimal value for F1 score resulted in similar performance for schema-1 and 3, but a balance of lower precision and higher recall for schema-2. Increasing the MQ threshold to 60 optimized precision but at considerable loss of recall for all three schemas (**Table 1**). Performance was most sensitive to the MQ threshold under schema 2 (no masking).

**Table 1.** Comparison of performance of proposed genome-masking schemas for SNS variant calling. For each masking scheme and MQ filtering threshold shown, the corresponding mean Precision, Recall, and F1 score is shown across all 36 Mtb isolates. Corresponding Precision-Recall curves are given in Figure 5A. Performance at a threshold of MQ≥40 is given as a common point of comparison across the three masking schemas.

| Masking Schema | Metric Optimized | MQ Threshold | F1 | Precision | Recall |
| --- | --- | --- | --- | --- | --- |
| Masking non-unique regions | F1-score | 19 | 0.87 | 99.77% | 77.98% |
|  | Comparator | 40 | 0.88 | 99.79% | 77.86% |
|  | Precision | 60 | 0.82 | 99.94% | 70.00% |
| No masking | F1-score | 8 | 0.94 | 98.56% | 88.95% |
|  | Comparator | 40 | 0.92 | 99.13% | 85.77% |
|  | Precision | 60 | 0.83 | 99.90% | 72.06% |
| Masking PLC genes (current standard) | F1-score | 35 | 0.82 | 99.50% | 70.30% |
|  | Comparator | 40 | 0.82 | 99.62% | 70.17% |
|  | Precision | 60 | 0.77 | 99.97% | 63.56% |

For INDELs (1-15 bp), precision was comparable to SNSs (96.2% - 100%, **Figure 6B**), while recall was lower (48.9% - 82.4%). At a threshold of MQ ≥ 40, we observed the following mean INDEL performances: 1) Masking non-unique regions, F1 = 0.83 (Precision = 98.2, Recall = 72.1%), 2) no masking of the genome, F1 = 0.89 (Precision = 98.9, Recall = 80.8%), 3) Masking PLC genes, F1 = 0.76 (Precision = 99.1%, Recall = 61.5%). Variant calling performance of short (1-5bp) INDELs was comparable to SNSs, and the limited performance for INDELs was largely driven by low recall of longer (6-15bp) INDELs (**Supp. Figure 5, Additional File 11**).

### Structural variation

We assessed the effect of structural variation (SV), of length ≥ 50 bp, a common source of reference bias, on variant calling performance (**Methods**). Detected SVs included the known regions of difference associated with Mtb Lineages 1, 2 and 3 (RD239, RD181, RD750 respectively)^25,26^ (**Supp. Figure 6**). Across all 36 isolate assemblies, we observed a strong negative correlation between average nucleotide identity to the H37Rv reference and the number of SVs detected (Spearman’s R = −0.899, p < 1.1e-13, **Supp. Figure 7)**. Additionally, we observe that 70% of detected SVs overlapped with regions with low pileup mappability (P-Map-K50E4 < 1.0).

We compared SNS variant calling performance by proximity to an SV and sequence uniqueness (**Figure 7, Additional File 12**), dividing variants into four groups: (1) SNSs in regions with perfect mappability (Pmap-K50E4 = 1) with no identified SV (87.3% of total 47,412 SNSs), (2) SNSs in regions with low mappability (Pmap-K50E4 < 1) with no identified SV (10.9% of SNSs), (3) SNSs in regions with perfect mappability within 100 bp of any identified SV (0.8% of SNSs), and (4) SNSs in regions with low mappability within 100bp of any identified SV (1.0% of SNSs). Variant calling performance decreased most sharply in regions with evidence for structural variation, especially when sequence content is also non-unique (Region types 3 & 4 respectively). Additionally, region type (2), or low mappability sequence content with no nearby SV, demonstrated reduced performance.

**Figure 7.**
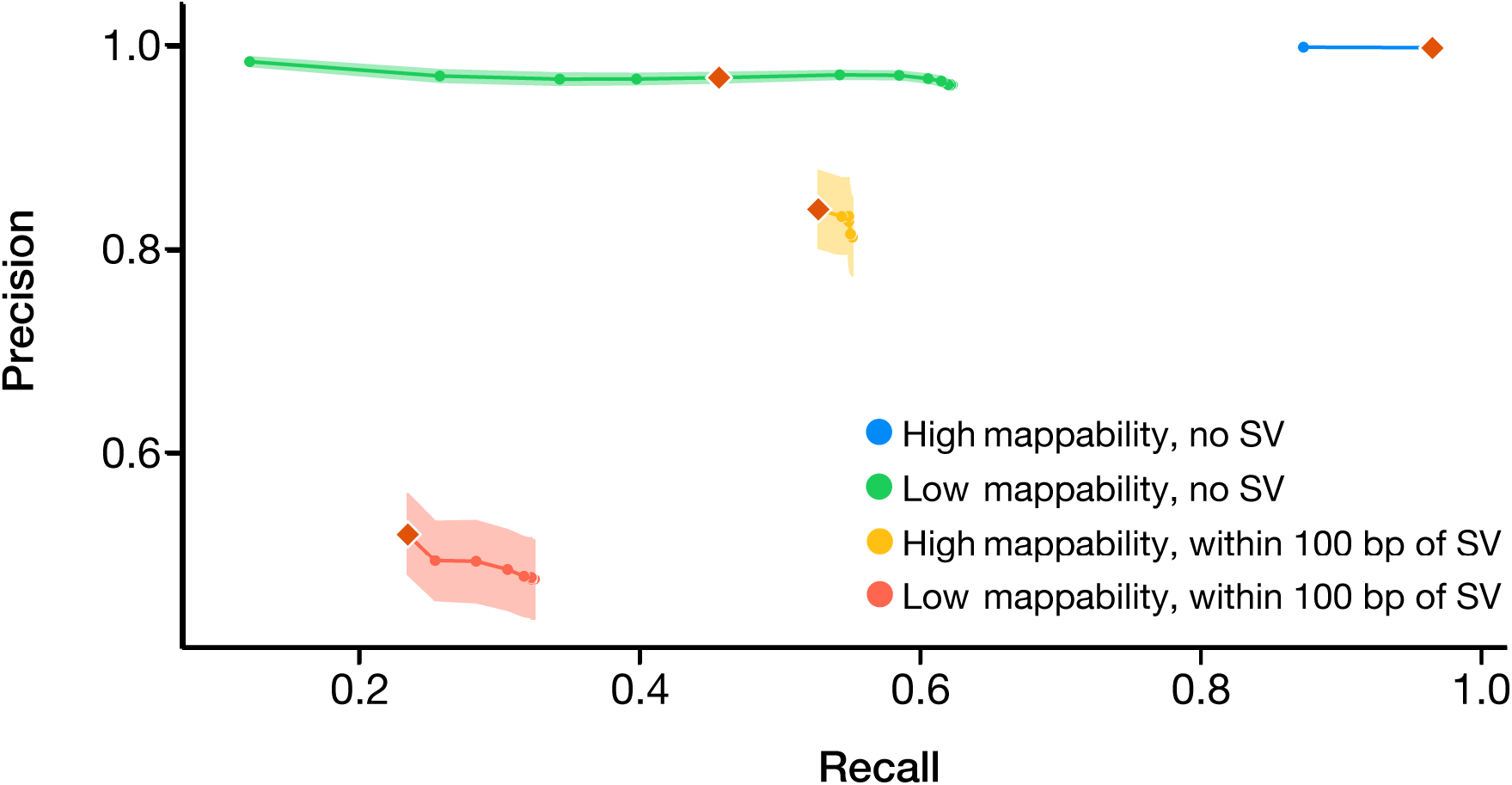
Variant calling performance of single nucleotide substitutions stratified by proximity to structural variants and low pileup mappability sequence. Mappability is dichotomized at Pmap-K50E4 =100% or <100%. Regions within 100bp of a SV categorized as “with SV”. Precision and recall is plotted for the following genomic contexts: (1) regions with high mappability with no SV (Blue, F1 = 0.98 (precision = 99.89%, recall = 96.49%, MQ threshold of 40)), (2) regions with low mappability and no SV (green, F1 = 0.62 (precision = 96.98%, recall = 45.65%, MQ threshold of 40), (3) regions with high mappability with SV (orange, F1 = 0.64 (precision = 84.07%, recall = 52.73%, MQ threshold of 40), (4) regions with low mappability and with SV (red, F1 = 0.32 (precision = 52.10%, recall = 23.47%, MQ threshold of 40). The standard error of the mean (SEM) for precision is shaded for each curve. Orange diamonds represent the precision and recall using the same MQ threshold of 40. Genomic contexts not near SVs (1 and 2) were evaluated with MQ thresholds ranging from 1-60. For genomic contexts within 100 bp of an SV (3 and 4), the MQ thresholds evaluated ranged from 1-40. Complete benchmarking results can be found for each individual isolate in Additional File 12.

### Refined regions of low confidence

Based on the presented analysis, we define a set of refined low confidence (RLC) regions of the Mtb reference genome. The RLC regions are defined to account for the largest sources of error and uncertainty in analysis of Illumina WGS, and is defined as the union of A) The 30 false positive hot spot regions identified (65 kb), B) low recall genomic regions with EBR < 0.9 (142 kb with 30 kb overlap with (A)), and C) regions ambiguously defined by long-read sequencing (**Methods**, 16 kb). We additionally evaluated the overlap between all detected SVs and the three RLC categories: RLC subset (A) overlapped 28% of SVs, RLC subset (B) overlapped with 65% of SVs, RLC subset (C) overlapped with 14% of SVs.

In total, the proposed RLC regions account for 177 kb (4.0%) of the total H37Rv genome (**Additional File 13**) and their masking represents a conservative approach to variant filtering. Across the 36 isolates evaluated, masking of the RLC regions combined with a SNS filter of MQ ≥ 40 would produce a mean F1-score of 0.882, with a mean precision of 99.9% and a mean recall of 78.9%.

## Discussion

The analysis and interpretation of Illumina WGS is critical for both research and clinical applications. Here, we study the ‘blindspots’ of paired-end Illumina WGS by benchmarking reference-based variant calling accuracy using 36 Mtb isolates with high confidence complete genome assemblies. Overall, our results improve our general understanding of the factors that affect Illumina WGS performance. In particular, we systematically quantify variant calling accuracy and the effect of sequence uniqueness, GC-content, coverage bias, and structural variation. For Mtb, we demonstrate that a much greater proportion of the genome can be analyzed with Illumina WGS than previously thought and provide a systematically defined set of low confidence/troublesome regions for future studies.

Approaches to benchmarking variant calling from Illumina WGS vary by field and species of interest and more standardization is needed^27^. Variant calling accuracy is usually benchmarked through *in silico* variant introduction with read simulation or otherwise using a small number of reference genomes that seldom capture the full range of diversity within a particular species. Our benchmarking exercise is unique in using a large and diverse set of high quality genome assemblies that are built using a hybrid long and short read approach. We further demonstrate that PacBio long-read sequencing is much less prone to coverage bias and is able to generate complete circular bacterial assemblies bridging repetitive regions in the majority of isolates with a median depth > 180×. The assemblies we generate will be an important community resource for benchmarking future variant calling or other WGS based bioinformatics tools.

The benchmarking results clearly demonstrate that low variant recall is a major limitation of reference-based Illumina variant calling, which achieved at most 89% recall at the optimal F1-score. Precision of variant calling using Illumina on the other hand was very high, with the small number of false variant calls concentrated in repetitive and structurally variable regions. We find that the best balance between precision and recall is achieved by tuning the variant mean mapping quality threshold, i.e. confidence of the read mapping. The specific mapping quality threshold will likely vary by species. For a GC-rich organism with highly repetitive sequence content like Mtb, a threshold of 40 achieved 85.8% recall and 99.1% precision.

Studying specific sources of low recall from Illumina, we identified insufficient read coverage to be the major driver, due not only to repetitive sequence content but also due to high-GC content and other sources of coverage bias. We further identified regions near structural variation to be particularly prone to low recall and precision. Of the variants we study, longer INDELs were recalled at lower rates than SNSs or INDELs < 6bp in length. These observations support ongoing efforts by the bioinformatics research community to build graph-reference genomes and align short reads to these graphs. Using a graph pan-genome built with a diverse set of Mtb reference genomes, there is great potential to both increase recall and precision of variant calling in divergent regions of the genome.

An alternative and generalizable approach to balancing precision and recall of reference-based Illumina variant calling is to mask repetitive (low mappability) regions. This simple approach does not require tuning the mapping quality threshold against a ground truth set of assemblies and relies instead on computing the pileup mappability metric across the reference sequence. This fills a gap for variant calling in other organisms using short-read mapping where low confidence regions may not already be defined. Compared with tuning against a ground-truth set of assemblies, this masking approach is conservative: for Mtb and filtering by MQ ≥ 40, precision is slightly higher at 99.8% vs 99.1% respectively and recall is lower at 77.9% vs 85.8% respectively.

Given Mtb’s genomic stability and clonality, this organism is particularly well suited for systematically identifying the sources of variant calling error from short-read data. Although 10.7% of the Mtb reference sequence is commonly excluded from genomic analysis, our results demonstrate that more than half of these regions are accurately called using Illumina WGS. For the PE/PPE family, of highest concern for sequencing error, nearly one third (52/168) had perfect mappability and near perfect gene-level EBR (≥ 0.99). The PE/PPE genes with poor performance were largely the PE_PGRS and PPE_MPTR sub-families. Only 65 kb (1.5%) of the reference genome H37Rv were responsible for the majority of false positives (89.2% of false positives across 36 isolates).

We present a set of refined low confidence (RLC) regions of the Mtb genome, designed to account for the largest sources of error and uncertainty in analysis of Illumina WGS (**Additional File 13**). Long-read data can allow RLC regions to be defined for other species to improve accuracy of Illumina WGS. The Mtb RLC regions span 4.0% of the reference genome, and their masking provides a conservative approach to variant calling, appropriate for applications where precision is prioritized over recall. At the same time, RLC region masking offers higher recall than the current field standard where more than 10% of the Mtb reference genome is masked. One limitation is that RLC regions were largely defined based on EBR of Illumina sequencing in our dataset that was restricted by design to 100+ bp paired end sequencing. We do not recommend the use of these RLC regions for Illumina sequencing at shorter read lengths or single-end reads. Instead we make available a more appropriate masking scheme of RLC regions + low pileup mappability (**Additional File 14**). Another limitation is that we defined RLC regions using the same set of high confidence assemblies evaluated. The reported precision and recall with RLC region masking are thus likely overestimates. On the other hand, we expect precision and recall estimates of the alternative approaches of masking low mappability regions or filtering at MQ ≥ 40 to be more robust.

Improving Illumina variant recall has significant implications. For clonal Mtb, for example, transmission inference using genomic data often relies on a very small number of SNS or INDEL differences between genome pairs. The observed large increase in recall we observe has the potential to substantially improve transmission inference^28^and/or our understanding of genome stability and adaptation.

## Conclusions

In summary, we show that Illumina whole genome sequencing has high precision but limited recall in repetitive and structurally variable regions when benchmarked against a diverse set of complete assemblies. We demonstrate that filtering variants using the mean mapping quality against a achieves the highest balance of precision and recall. Masking repetitive sequence content is a second generalizable solution, albeit a more conservative one, that maintains high precision. For Mtb, these two approaches increase recall of variants by 15.6% and 7.7% respectively, with a minimal change in precision (−0.5% and +0.1% respectively at MQ ≥ 40), allowing high variant recall in >50% of regions previously considered by the field to be error-prone. Our results improve variant recall from Illumina data with broad implications for clinical and research applications of sequencing. We also provide a high-quality set of genome assemblies for benchmarking future variant calling or other WGS based bioinformatics tools.

## Methods

### Summary of sequencing data used

Our dataset consisted of a convenience set of 16 clinical isolates from Lima, Peru, previously sequenced with Illumina WGS and archived in frozen culture^29^. These isolates were revived and sequenced with PacBio RS II long-read sequencing (Dataset #1). Additionally, 15 total clinical isolates isolated in Azerbaijan, Georgia, Moldova were sequenced with PacBio Sequel II long-read sequencing^30^(Dataset #2).

This dataset of 31 clinical isolates was combined with publicly available paired PacBio (RS II) and Illumina genome sequencing from 19 clinical isolates from two previously published studies^20,21^. From these four sources, 38 Mtb isolates were selected for having a) Illumina WGS with paired end reads with at least a median sequencing depth of 40X relative to the Mtb reference genome (H37Rv). All aggregated metadata and SRA/ENA accessions for PacBio and Illumina sequencing data associated with this analysis can be found in **Additional File 15**.

### DNA extraction for PacBio (RS II) Sequencing of Peruvian Isolates (Data Source #1)

MTB cultures were allowed to grow for 4-6 weeks. Pellets were heat-killed at 80°C for 20 minutes67,68, the supernatants were removed, and the enriched cell pellet was subjected to DNA extraction soon after or stored frozen until extraction. Largely intact DNA was extracted from heat-killed cells pellets using a protocol tailored for mycobacteria that ends with a column-based elution^31^. Yields were determined using fluorescent quantitation (Qubit, Invitrogen/Thermo Fisher Scientific) and quality was assessed on a 0.8% GelRed agarose gel with 1XTAE, separated for 90 minutes at 80V.

### PacBio (RS II) Sequencing of Peruvian Mtb Isolates (Data Source #1)

Approximately 1 μg of high molecular weight genomic DNA was used as input for SMRTbell preparation, according to the manufacturer’s specifications (SMRTbell Template Preparation Kit 1.0, Pacific Biosciences). Briefly, HMW gDNA was sheared to 20kb using the Covaris g-tube at 4500 rpm. Following shearing, gDNA underwent DNA damage repair, ligation to SMRTbell adaptors and exonuclease treatment to remove any unligated gDNA. At least 500 ng final SMRTbell library per sample was cleaned with AMPure PB beads and 3-50 kb fragments were size selected using the BluePippin system on 0.75% agarose cassettes and S1 ladder, as specified by the manufacturer (Sage Science). Size selected SMRTbell libraries were annealed to sequencing primer and bound to the P6 polymerase prior to loading on the RSII sequencing system (Pacific Biosciences). Sequencing was performed using C4 chemistry and 240-minute movies. Following data collection, raw data was converted into subreads for subsequent analysis using the RS_Subreads.1 pipeline within SMRTPortal (version 2.3), the web-based bioinformatics suite for analysis of RSII data.

### DNA extraction for PacBio (Sequel II) Sequencing (Data Source #2)

For all samples from Azerbaijan and Georgia, MTB cultures were grown in 7H9+ADST broth to A600 0.5–1.0. Pelleted cells were heat killed at 80°C for 2 hours. Cell pellets were resuspended in 450ul TE-Glu, 50ul of 10 mg/mL lysozyme was added and incubated at 37°C overnight. To each sample 100ul of 10% sodium dodecyl sulfate and 50ul of 10 mg/ml proteinase K was added and incubated at 55°C for 30 minutes. 200 ul of 5M sodium chloride and 160 ul Cetramide Saline Solution (preheated 65°C) was added then incubated for 65°C for 10 minutes. To each sample 1 ml chloroform:isoamyl alcohol (24:1) was added, mixed gently by inversion. Samples were centrifuged at 5000g for minutes, and 900ul of aqueous layer was transferred to fresh tube. DNA was re-extracted with chloroform:isoamyl alcohol (24:1) and 800 ul of aqueous layer was transferred to fresh tube. To 800 aqueous layer 560 ul isopropanol was added, mix gently by inversion. The precipitated DNA was collected by centrifuging for 10 minutes and supernatant was removed. DNA was washed with 70% ethanol, and DNA was collected by centrifuging and supernatant removed. Air dried DNA pellet was dissolved overnight in 100 ul of TE buffer, and stored at 4°C.

For all samples from Moldova, DNA was extracted according to CTAB protocol^32^.

### PacBio (Sequel II) Sequencing (Data Source #2)

Approximately 1 μg of high molecular weight genomic DNA was used as input for SMRTbell preparation according to the manufacturer’s protocol (Preparing Multiplexed Microbial Libraries Using SMRTbell Express Template Prep Kit 2.0, Pacific Biosciences). Briefly, HMW gDNA was sheared to ~15kb using the Covaris g-tube at 2029 × g. For about half of the samples the molecular weight of the DNA did not need shearing. Following shearing, gDNA underwent DNA damage repair, ligation to SMRTbell barcoded adaptors and exonuclease treatment to remove any unligated gDNA. At least 500 ng of pooled SMRTbell library per sample was cleaned with AMPure PB beads and 7-50 kb fragments were size selected using the BluePippin system on 0.75% agarose cassettes and S1 ladder, as specified by the manufacturer (Sage Science). The pool of size-selected SMRTbell libraries were annealed to v4 sequencing primer and bound to the polymerase prior to loading on the Sequel II sequencing system (Pacific Biosciences). Sequencing was performed using version 1 chemistry and 15-hour movies.

### H37Rv reference genome and gene annotations

The H37Rv (NCBI Accession: NC_000962.3) genome sequence and annotations was used as the standard reference genome for all analyses. Functional category annotations for all genes of H37Rv were downloaded from Release 3 (2018-06-05) of MycoBrowser^33^(https://mycobrowser.epfl.ch/releases). PE/PPE sub-family annotations of H37Rv were taken from Ates et al.^34^. Programmatic visualization of data along with annotations of the H37Rv genome were made using the DNA Features Viewer python library^35^.

### Genome assembly with PacBio long-read data

All PacBio reads were assembled using Flye^36^(v2.6). After assembly, Flye performed three rounds of iterative polishing of the genome assembly with the PacBio subreads, producing a polished de novo PacBio assembly. If Flye identified the presence of a complete circular contig, Circlator^37^(v1.5.5) was used to standardize the start each assembly at the DnaA (Rv0001) locus.

### Polishing of *de novo* PacBio assemblies with Illumina WGS

The paired-end Illumina WGS reads were trimmed with Trimmomatic^38^(v0.39) with the following parameters: 2:30:10:2:true SLIDINGWINDOW:4:20 MINLEN:75. Trimmed reads were aligned to the associated de novo PacBio assembly with BWA-MEM^39^(v0.7.17). Duplicate reads were removed from the resulting alignments using PICARD^40^ (v2.22.5). Using the deduplicated alignments, Pilon^22^(v1.23) was then used to correct SNSs and small INDELs in the *de novo* PacBio assembly, producing a high confidence assembly polished by both PacBio and Illumina WGS.

### Identifying mixed infections using F2 metric and removing mismatched PacBio and Illumina WGS

To further reduce the effects of contamination, we used the F2 metric to identify samples that may have inter-lineage variation due to co-infection^<u>41</u>^. The F2 metric measures the heterogeneity of genotypes at known lineage defining positions of the H37Rv genome. We computed the F2 lineage-mixture metric for both PacBio and Illumina WGS from each isolate. Isolates were filtered out if either the F2 metric for Illumina sequencing passed 0.05 or the F2 metric for PacBio sequencing passed 0.35. The threshold used for PacBio sequencing subreads is much higher because the inherent error rate per read is much higher than Illumina.

During polishing we identified the N0052 isolate from Chiner-Oms et al.^20^as a potential sample mismatch, meaning PacBio and Illumina WGS were not performed on the same clinical isolate. When polishing the de novo assembly of N0052, we found that the following changes were performed based on the Illumina WGS: 594 SNPs, 19 insertions, and 92 deletions. The extreme number of corrected SNPs by Illumina polishing is drastically different from the known error profile (**Additional File 2-3**). Additionally, the inferred sub-lineage of the de novo PacBio assembly was lineage 2.2.1, while the inferred sub-lineage based on Illumina WGS and the Illumina Polished PacBio assembly was lineage 2.2.2 (**Additional File 2**). The fact that the polishing with Illumina WGS changed known lineage defining SNPs makes the sample further suspect as a mismatch. Thus, N0052 was removed from analysis as to minimize chances of benchmarking wrongly matched data.

### Evaluation of PacBio genome assembly characteristics and multiple genome alignment

FastANI^42^was used to calculate the average nucleotide identity to the H37Rv reference genome for all completed genome assemblies. The Prokka (v1.13) genome annotation pipeline^43^ was used to annotate genes in each completed genome assembly. The genome size and GC content of the entire genome was calculated from each assembly using custom python code. The progressiveMauve algorithm of the Mauve (v2.4.0)^44^alignment software was used to perform multiple sequence alignment of all 36 completed Mtb assemblies and the H37Rv reference genome (NCBI Accession: NC_000962.3). The multiple genome alignments of H37Rv and 36 assemblies were visualized using the Mauve GUI^45^(**Supp. Figure 2**).

### Variant calling and structural variant detection using complete PacBio assemblies

Minimap2^46^was used to align each polished circular completed assembly to the H37Rv reference genome, producing a base-level alignment of similar regions of the assembly to H37Rv. In regions with high sequence diversity or large structural variation, Minimap2 will not produce alignments. To account for this, the NucDiff^47^analysis pipeline, which uses the MUMmer^48^aligner internally, was also used to detect and classify the presence of large structural variants relative to the H37Rv reference. All structural variants (≥ 50 bp) identified by NucDiff for each genome assembly can be found in (**Additional File 16**).

### Illumina WGS data processing for variant calling relative to H37Rv

Paired-end Illumina reads were trimmed with Trimmomatic (v0.39) with the following parameters: 2:30:10:2:true SLIDINGWINDOW:4:20 MINLEN:75. Trimmed reads were aligned to the H37Rv reference genome (NC_000962.3) with BWA-MEM^39^ (v0.7.17). Duplicate reads were removed from the resulting alignments using PICARD^40^(v2.22.5). Using the deduplicated alignments, small genome variants (SNSs and INDELs) were inferred using Pilon^22^(v1.23). Samtools, Bcftools, and BEDtools were used as needed for SAM/BAM, and VCF/BCF format file manipulation^49–51^.

### Phylogenetic inference using complete genome assemblies

All single nucleotide variants inferred through alignment with Minimap2 of PacBio assembly to the H37Rv genome were concatenated across the 36 strains. Any SNS position which was ever ambiguously called in at least 1 isolate was excluded (No NAs allowed, only REF or ALT alleles allowed). Thus, in order for a SNS position to be included it needed to have no ambiguity relative to the H37Rv reference in any isolate. FastTree^52^was used to infer an approximate maximum likelihood phylogeny from the concatenated SNS alignment of all 36 clinical Mtb isolates (15,673 total positions across 36 Mtb clinical isolates).

### Measuring repetitive sequence content of the H37Rv reference genome using Pileup Mappability

We evaluated sequence uniqueness using a *mappability* metric defined as the inverse of the number of times a sequence of length *K* appears in a genome allowing for *e* mismatches and considering the reverse complement^53^. The *pileup mappability* of a position in a genome is then defined as the average mappability of all overlapping k-mers. Thus, there are 2 parameters when calculating mappability, k (length of k-mer) and e (number of base mismatches allowed in counting matching k-mers). Genmap^54^(v1.3) was used to calculate the mappability of all k-mers across the H37Rv reference genome with the following parameters: k-mer sizes of 50, 75, 100, 125, 150 base pairs and E = 0-4 mismatches. The Gene-level mappability (k = 50 bp, e = 4 mismatches) scores were computed as the average pileup mappability across all genes bodies annotated in H37Rv (NCBI Accession: NC_000962.3). The base level pileup mappability scores of H37Rv are available in TSV and BEDGRAPH format for easy visualization in a genome browser (**Additional Files 6 and 17**).

### Calculation of Empirical Base-level Recall (EBR) of Illumina variant calling

The goal of the empirical base-level recall (EBR) for score is to summarize the consistency by which Illumina WGS correctly evaluated any given genomic position. The EBR for a genomic position was defined as the proportion isolates where Illumina WGS confidently and correctly agreed with the PacBio defined ground truth. The ground truth was inferred for each isolate by directly comparing the completed PacBio genome assembly to the H37Rv reference using Minimap2^46^and NucDiff^47^. Due to Minimap2’s inability to classify large structural variants, the ground truth relative to H37Rv was supplemented with the structural variant calls generated by the NucDiff analysis pipeline. Illumina WGS reads were aligned to the H37Rv reference genome with BWA-MEM^39^, and variants were inferred with the Pilon^22^ variant detection tool. In addition to identifying variants relative to the reference genome, Pilon provides variant calling annotations for all positions of H37Rv. The variant calling quality annotations of Pilon for all positions of H37Rv were parsed for comparison to the PacBio defined ground truth for each isolate evaluated.

Only the following comparison outcomes were classified as a correctly recalled position:

1) Both Illumina variant calling and the PacBio ground truth agree on the genotype of a genomic position, 2) Both Illumina variant calling and the PacBio ground truth agree that a genomic position is deleted.

The following comparison outcomes were classified as poorly recalled position:

3) The PacBio ground truth supports a deletion, but Illumina is not confident in the presence of the deletion, 4) Both Illumina variant calling and the PacBio ground truth disagree on the genotype of a genomic position, 5) The PacBio ground truth supports the presence of a genomic region, while Illumina variant calling did not confidently support the presence of the region. 6) Illumina variant calling erroneously supports a deletion at a genomic position which is not deleted in the PacBio ground truth.

The following EBR comparison outcomes were classified as ambiguous (N/A) due to ambiguities in the interpretation of the ground truth: a) Cases where the PacBio ground truth contained genome duplications relative to H37Rv, b) Cases where the PacBio ground truth did not provide a confident alignment or structural variant call due to high sequence divergence from the reference sequence.

For calculating the EBR for a genomic position, ambiguous (N/A) outcomes were ignored when the number of N/As was <= 25%. In the case that a position had greater than 25% N/As at a genomic position, the EBR score was defined as “Ambiguous”. Ambiguous (N/A) EBR scores represent locations of the H37Rv genome where there appeared to be systematic trouble in determining the ground truth genotype.

The base level EBR scores are available in TSV and BEDGRAPH format for easy visualization in a genome browser (**Additional Files 6 and 18**).

### Evaluating characteristics of low empirical performance across Mtb genome

The Illumina WGS variant caller used, Pilon, produces VCF tags for all reference positions evaluated, including positions which were confidently called a reference. The tags associated with each position can either be PASS or a combination of non-pass tags (LowCov, Del, Amb). Each genomic position can be assigned a combination of the following VCF Tags: a) PASS, signifying confirmation of either a reference or an alternative allele. b) LowCov, signifying insufficient high confidence reads (Depth < 5). c) Del, signifying that the position is confidently inferred to be deleted. d) Amb, signifying evidence for more than one allele at this position. We quantified the frequency of all combinations of these tags across all positions that were classified as “poor recalled” during EBR evaluation.

### Measuring sequencing bias with per-base relative depth

We measured sequencing bias using the relative depth statistic, which for a given genome assembly and sequencing dataset, is defined as the sequencing depth per site divided by average depth across the entire genome^4^. We evaluated the relative depth of all base pair positions of all sequencing runs (Illumina and PacBio) relative to the corresponding isolates’ complete PacBio genome assembly. The sequencing depth of a base pair position was defined as the number of reads with a nucleotide aligning to the position of interest. We calculated the mean coverage across a sample by simply averaging the depth across all positions of the evaluated genome. For ambiguous mapping reads, the aligners used (BWA-mem and Minimap2) use a random assignment policy between all possible alignment locations. This allows for approximation of depth in regions with non-uniquely mapping reads. For each individual Mtb isolate, we then calculated the mean relative depth across all positions with the same GC content (100 bp window size, **Additional File 8**).

### Defining and excluding ambiguous regions relative to H37Rv (per isolate genome assembly)

Following GA4GH (Global Alliance for Genomics & Health) benchmarking guidelines^23^, we excluded regions of the genome, where definition of the ground truth had ambiguity in its definition relative to the reference genome. The following comparison outcomes were classified as ambiguous (N/A) due to ambiguities in the interpretation of the ground truth: a) Cases where the PacBio ground truth contained duplications relative to H37Rv, b) Cases where the PacBio ground truth did not provide a confident alignment or structural variant call due to high sequence divergence relative to H37Rv. These regions thus represent sequences of divergence relative to the reference genome.

The percentage of the reference genome that was identified as “ambiguous” was consistently less than 1% for all 36 clinical isolates evaluated. The median percent of the genome where the ground truth was “ambiguously defined” was 0.4% (IQR: 0.3% − 0.5%). A large majority of these ambiguous ground truth regions were either in Mobile Genetic Elements, PE_PGRS or PPE_MPTR genes. The ambiguously defined regions for each isolate can be found in **Additional File 4**. Additionally, all regions of the H37Rv genome which were ambiguous in over 25% of isolates, signifying high levels of ambiguity, are present in **Additional File 5**.

### Defining the putative low confidence (PLC) regions of the H37Rv genome

The regions most commonly excluded from Mtb genomics analysis, also referred to as the Putative Low Confidence (PLC) regions in this work, were based on current literature^16,24,55,56^. Specifically, we defined the PLC regions as the union of the 168 PE/PPE genes, all mobile genetic elements (MGEs), and 82 genes with repetitive content previously identified^24^. PLC regions are defined in **Additional File 19** (BED format). Non-PLC regions were simply defined as the complement of the PLC genes.

### Evaluating variant calling performance of genome masking approaches

Following the small variant benchmarking standards outlined by the GA4GH, we used Hap.py (v0.3.13) to evaluate the Illumina WGS variant calling performance of Pilon for all 36 isolates individually. For each complete genome assembly, SNSs and small INDELs 1-15 bp inferred by the Minimap2-paftools pipeline were used as ground truth. We evaluated variant calling performance of Illumina WGS when using different region filtering schemas: (1) masking of all PLC genes, the current standard practice, (2) masking of repetitive regions with P-Map-K50E4 < 100%, and (3) No masking. Masking schemas (1 and 2) are provided in BED format (**Additional File 19 and 20**). After applying each masking schema, we filtered potential variants according to whether the Pilon variant calling pipeline gave the variant a PASS filter and the mean mapping quality (MQ) of all reads aligned to the variant position.

For each combination of region masking and variant filtering using mapping quality, we then calculated the absolute number of true positives (TP, i.e. a variant in the ground truth variant set and correctly called by the Illumina variant calling pipeline), false positives (FP, the Illumina variant calling pipeline calls a variant not in the ground truth set), and false negative (FN, the variant is in the ground truth set but is not called by the Illumina variant calling pipeline) variant calls. For each set of parameters, we calculated the overall precision (positive predictive value) as TP/(TP + FP), and recall (sensitivity) as TP/(TP + FN). In agreement with the default behavior of Hap.py, and to avoid undefined precision values, filtering parameters that yielded no TP or FP were defined as having a precision of 1.0 and a recall of 0. Additionally, we calculated the F1-score, which weights precision and recall with equally: F1 = 2 * (precision * recall)/(precision + recall). The F1 score summarizes each variant calling performance as a single value between 0 and 1 (where 1 represents both perfect precision and recall).

To aggregate the performance evaluation across all 36 isolates, the mean and standard error of the mean (SEM) of precision, recall and F1 score was calculated for all sets of parameters evaluated (**Additional File 10**). The individual variant calling performance statistics for each isolate can also be found in **Additional File 10**. The variant calling performance comparison of shorter (1-5bp) vs longer (6-15bp) INDELs can be found in **Additional File 11**.

### Evaluating variant calling performance near regions with structural variation and repetitive sequence content

Using Hap.py and the same approach defined in the above section, we evaluated SNS variant calling performance in the following types of regions: (1) SNSs in regions with perfect mappability (Pmap-K50E4 = 1) with no identified SV (2) SNSs in regions with low mappability (Pmap-K50E4 < 1) with no identified SV, (3) SNSs in regions with perfect mappability within 100 bp of any identified SV, and (4) SNSs in regions with low mappability within 100bp of any identified SV. Genomic contexts not near SVs (1 and 2) were evaluated with MQ thresholds ranging from 1-60. For genomic contexts within 100 bp of an SV (3 and 4), the MQ thresholds evaluated ranged from 1-40. The MQ threshold evaluated near SVs was limited due to the fact that a majority of SNSs near SVs typically have lower MQ values, and higher MQ values resulted in recalls of approximately As explained in the previous section, the mean and SEM of precision, recall, and F1 score were calculated for all MQ filtering thresholds across all 4 region types (**Additional File 12**).

### Evaluation of the distribution of potential false positive SNS calls across the Mtb genome

False positive SNS calls were identified by the Hap.py evaluation software through comparison to the assembly-based ground truth variant call set. Additionally, false positive calls with MQ < 30 were filtered out, as to only include false positives which would realistically pass standard filtering. For each genomic region (gene or intergenic region) of the H37Rv genome, the total number of overlapping false positives across all 36 isolates was calculated (**Additional File 9**). Across all 36 clinical isolates, there were 548 false positive SNSs with MQ ≥ 30 and 696 total false positive SNS with MQ ≥ 1 detected.

### Defining Refined Low Confidence (RLC) regions

We defined the refined low confidence regions (RLC) of the Mtb reference genome as the union of A) The 30 false positive hot spot regions (gene and intergenic) identified (65 kb), B) poorly recalled genomic regions as identified by EBR (EBR < 0.9, 142 kb), and C) regions with frequently ambiguously defined ground truths (16 kb). We provide the complete set of RLC regions in BED format (177 kb, **Additional File 13**), along with each separate component of the RLC regions in BED format (**Additional Files 21, 22, and 23**). For very conservative masking of the Mtb reference genome, we additionally provide a masking scheme that specifies the union of a) the RLC regions and b) all low pileup mappability regions (PmapK50E4 < 1) (277 kb, **Additional File 14**).

## Author Contributions

MGM and MRF conceived, designed and conducted the study. MGM and MRF wrote the manuscript with input from all authors. RVJ provided bioinformatics support and input on data analysis. LEE, DD, M. Salfinger and M. Strong cultured Mtb isolates and performed DNA extraction in preparation for PacBio sequencing of Dataset #1. IA, SV, and VC cultured Mtb isolates and performed DNA extraction in preparation for PacBio sequencing of Dataset #2.. AR, MH, and BJ selected clinical isolates and assisted in data processing for PacBio sequencing of Dataset #2. ZI provided help and advice throughout the project. The final manuscript was read and approved by all authors.

## Data availability and materials

All new sequencing data generated for this study and complete Mtb genome assemblies were submitted to NCBI SRA and Genbank databases under BioProject accession number PRJNA719670 (Submission Pending). The publicly available PacBio and Illumina data from two previously published studies^20,21,57^ is available from PRJEB8783, PRJEB31443, PRJEB27802, and PRJNA598991. SRA/ENA accessions and related sequencing metadata for all data can be found in Additional File 15. All code for data processing and analysis in this study is available from the following GitHub repository, https://github.com/farhat-lab/mtb-illumina-wgs-evaluation. The repository README provides instructions to run each part of the analysis using the Snakemake^58^ workflow engine and using Python based Jupyter notebooks.

## Acknowledgments

We are grateful to Natalia Quiñones, and Karel Brinda for their helpful discussions and advise throughout the project. We grateful to Melissa Smith and Irina Oussenko for their assistance in PacBio (RS II) long read sequencing of the M. tuberculosis genomic DNA. We acknowledge NIH Intramural Sequencing Center (NISC) for the PacBio (Sequel II) long-read sequencing of the M. tuberculosis genomic DNA; Critical Path Institute (C-Path) and Translational Genomics Research Institute (T-Gen) for the Illumina sequencing of the M. tuberculosis DNAs and for the mTB DNA long-term storage; the International Science and Technology Center for their support in establishing the TB Portal agreement with Georgia; CRDF Global for their support in establishing the TB Portal agreements with Azerbaijan and Moldova. This research was supported in part by the Office of Science Management and Operations of the NIAID.

## Supplementary Information

Additional File 1: Supplementary Figures and Tables (Figure S1-7, Table S1-6)

Additional File 2: Results and quality control for assembly and sequencing for both PacBio and Illumina sequencing

Additional File 3: List of all changes made during Illumina polishing of the d*e novo* PacBio assemblies

Additional File 4: List of genomic regions with ambiguously defined ground truths relative to H37Rv for all each isolate evaluation

Additional File 5: List of genomic regions which were frequently had an ambiguously defined ground truth

Additional File 6: Table containing the EBR, Pileup Mappability, and GC% of all genomic positions of the H37Rv reference. Due to large file size, Additional File 6^59^ is hosted on Zenodo at https://zenodo.org/record/4662193.

Additional File 7: EBR, and Pileup Mappability across all genomic regions of H37Rv (both genes and intergenic regions)

Additional File 8: Table of the mean relative sequencing depth of both Illumina and PacBio at varying GC% across all 36 isolates evaluated.

Additional File 9: Table containing the frequency of observed False Positive SNSs (MQ ≥ 30) across all genomic regions of H37Rv (both genes and intergenic regions)

Additional File 10: Variant call benchmarking of SNSs and small indels (<=15bp)

Additional File 11: Variant call benchmarking stratified by shorter (< 6bp) and longer indels (6-15bp)

Additional File 12: Variant call benchmarking of SNSs stratified by proximity to an SV and low pileup mappability

Additional File 13: Masking scheme in BED format specifying the Refined Low Confidence Regions

Additional File 14: Masking scheme in BED format specifying the union of a) Refined Low Confidence Regions, and b) regions with Pileup Mappability (K= 50 bp, E = 4 mismatches) < 1.

Additional File 15: SRA/ENA sequencing run metadata for PacBio and Illumina sequencing used in this study

Additional File 16: All identified structural variants for each complete genome assembly as identified by the NucDiff analysis pipeline.

Additional File 17: Base-level Pileup Mappability scores (P-Map-K50E4) across the H37Rv in BEDGRAPH format

Additional File 18: Base-level EBR scores (36 isolates) across the H37Rv in BEDGRAPH format

Additional File 19: Masking scheme for the Putative Low Confidence (PLC) Regions in BED format

Additional File 20: All regions with low pileup mappability (P-Map-K50E4 < 100%) in BED format

Additional File 21: Component (A) of RLC regions. Masking scheme Specifying the 30 false positive hot spot regions (gene and intergenic) in BED format.

Additional File 22: Component (B) of RLC regions. Masking scheme specifying poorly recalled genomic regions as identified by EBR< 0.9) in BED format.

Additional File 23: Component (C) of RLC regions. Masking scheme specifying regions that frequently (> 25%) had an ambiguously defined ground truth in BED format. Same information as Additional File 5 but this file is instead in BED format.

