## Additional File 1, Supplemental Figures & Tables for "Genomic sequence characteristics and the empiric accuracy of short-read sequencing"

#### Supplementary Figures & Tables

Supplemental Figure 1.

A.

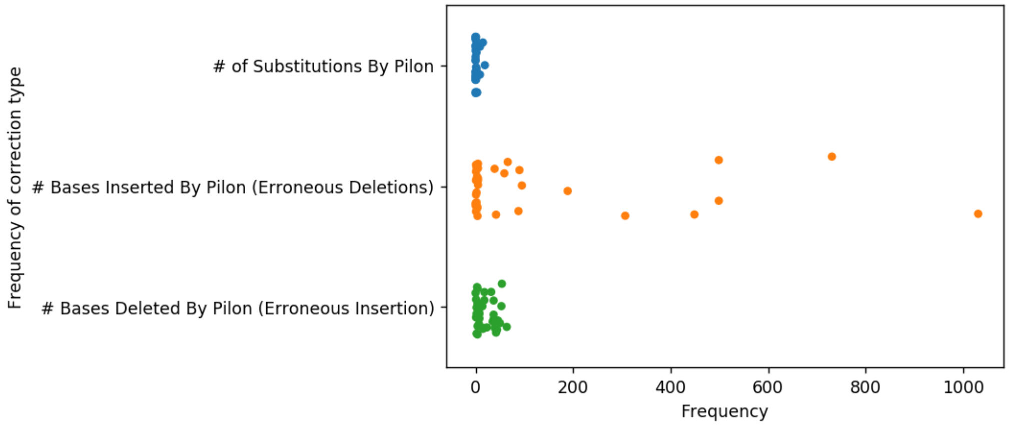


B.

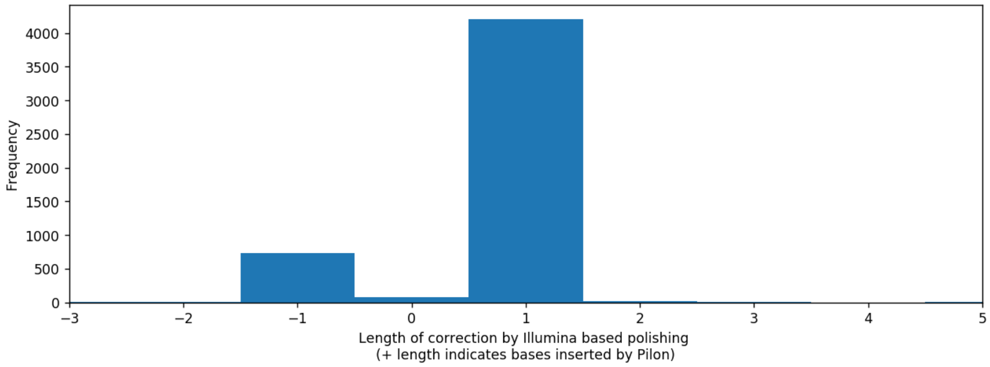


C.

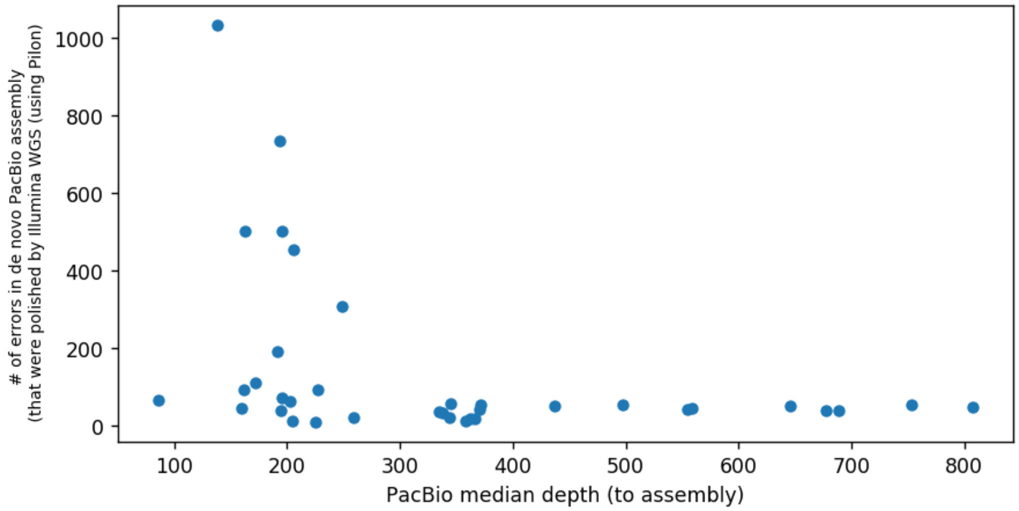

**Evaluation of profile of errors in *de novo* PacBio assemblies corrected by Illumina polishing.** a) The distribution of the number of types of corrections implemented by Illumina polishing of the de novo PacBio assemblies for each assembly. b) the distribution of nucleotide lengths of corrected made by Pilon during polishing. The insertion of bases (positive length) by Pilon indicates an erroneous deletion in the *de novo* PacBio genome assembly. c) We observed that the frequency of corrections (changes due to polishing) correlates with PacBio sequencing depth. We observed that the high coverage PacBio sequencing runs typically had a lower frequency of corrections made by Illumina WGS polishing with Pilon, while lower sequencing depth samples had higher numbers of erroneous deletions (Spearman's $R = -0.458$, $p < 4.9e-3$).

Supplemental Figure 2.

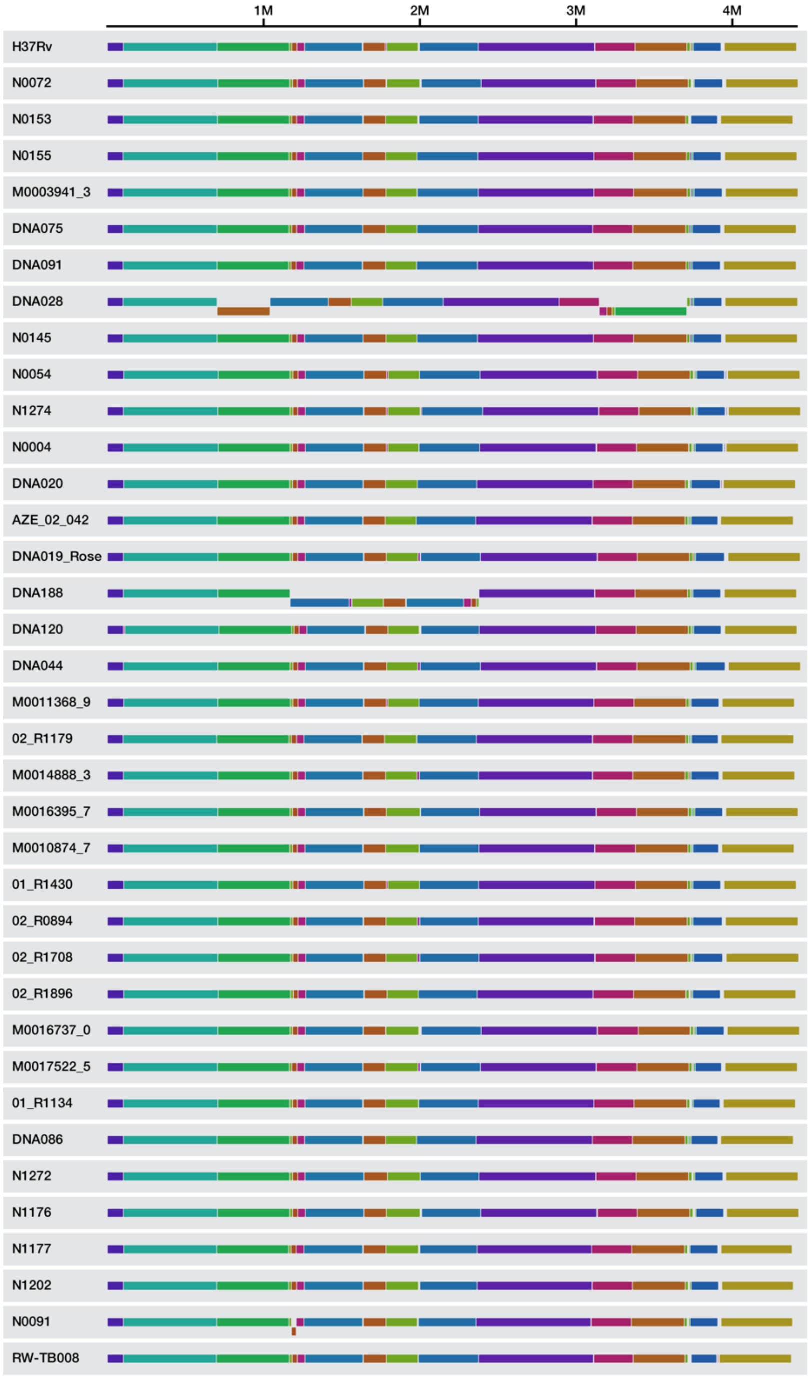


**Whole genome sequence alignment between the H37Rv reference genome and all 36 completed circular Mtb genome assemblies.** The whole genome multiple sequence alignment was performed using the *progressiveMauve*[44] algorithm. Each contiguously colored region is a locally collinear block (LCB), a region without rearrangement of homologous backbone sequence. LCBs below a genome's x-axis line represent a genome inversion that is in the reverse complement orientation relative to the reference genome, as observed in isolates N0091, DNA188, and DNA028.

#### Supplementary Table 1

| Region | Total bp | Mean EBR score | Median EBR Score | % of positions w/ EBR = 100% | % of positions w/ EBR ≥ 97% |
|---|---|---|---|---|---|
| Putative Low Confidence (PLC) Regions | 469,501 bp | 0.905 | 1.0 (IQR 0.917 - 1.0) | 59.8% | 66.8% |
| Non - PLC | 3,942,031 bp | 0.998 | 1.0 (IQR 1.0 - 1.0) | 97.6% | 99.3% |

**Information regarding the base-level distribution of EBR scores in PLC and non-PLC regions.**

#### Supplementary Table 2.

| Gene set | # genes | Median (+ IQR) mean-EBR score | # genes w/ mean-EBR < 97% |
|---|---|---|---|
| Non-PLC genes | 3695 | 1.0 (IQR: 0.9999 - 1.0000) | 14 (0.4%) |
| *pe/ppe* (PLC) | 168 | 0.9953 (IQR: 0.9026 - 0.9998) | 65 (38.7%) |
| Mobile genetic elements (PLC) | 147 | 0.9974 (IQR: 0.7647 - 1.0000) | 55 (37.4%) |
| Additional 69 repetitive genes* (PLC) | 69 | 0.9974 (IQR: 0.9859 - 0.9996) | 12 (17.4%) |

**Statistics from distribution of gene-level mean EBR between PLC and other regions.**

### Supplemental Figure 3.

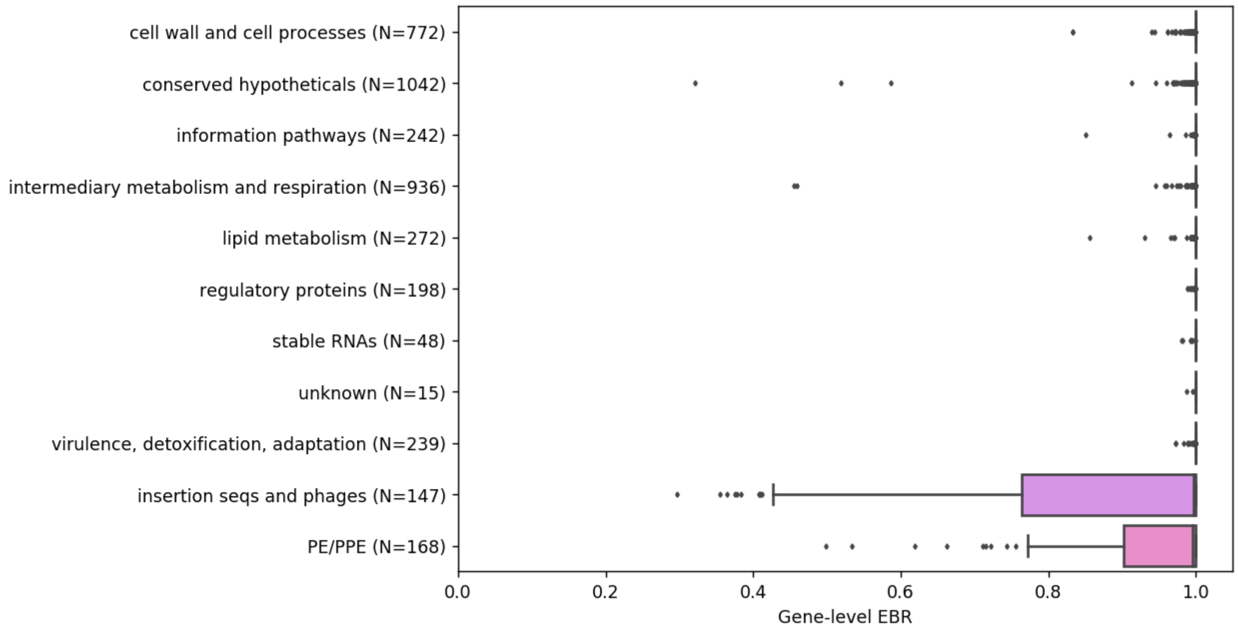


**The distribution of gene-level mean EBR across all annotated functional gene categories in the Mtb genome.** We compared the distribution of gene-level EBR scores across all functional gene categories annotated by MycoBrowser. The pe and ppe gene families (PE/PPEs) and mobile genetic elements (MGE), which make up 82% of PLC genes, demonstrated significantly lower mean EBR and Pileup Mappability than other non-PLC genes.

Supplemental Table 3.

| Gene set | # genes | Median (+ IQR) mean-EBR score | # genes w/ mean-EBR < 97% |
|---|---|---|---|
| cell wall and cell processes | 772 | 1.0 (IQR: 0.9997 - 1.0000) | 7 (0.9%) |
| conserved hypotheticals | 1042 | 1.0 (IQR: 1.0000 - 1.0000) | 7 (0.7%) |
| information pathways | 242 | 1.0 (IQR: 0.9999 - 1.0000) | 2 (0.8%) |
| intermediary metabolism and respiration | 936 | 1.0 (IQR: 0.9998 - 1.0000) | 6 (0.6%) |
| lipid metabolism | 272 | 1.0 (IQR: 0.9998 - 1.0000) | 4 (1.4%) |
| regulatory proteins | 198 | 1.0 (IQR: 1.0000 - 1.0000) | 0 (0%) |
| unknown | 15 | 1.0 (IQR: 1.0000 - 1.0000) | 0 (0%) |
| virulence, detoxification, adaptation | 239 | 1.0 (IQR: 1.0000 - 1.0000) | 0 (0%) |
| stable RNAs | 48 | 1.0 (IQR: 1.0000 - 1.0000) | 0 (0%) |
| *PE/PPE* (Part of PLC regions) | 168 | 0.9953 (IQR: 0.9026 - 0.9998) | 65 (38.7%) |
| Mobile genetic elements (Part of PLC regions) | 147 | 0.9974 (IQR: 0.7647 - 1.0000) | 55 (37.4%) |

**Statistics from distribution of gene-level mean EBR across all annotated functional gene categories in the Mtb genome.**

#### Supplementary Table 4.

| **Pilon Variant Calling Tag(s)** | Total Low Recall Positions (across all 36 isolates evaluated) | % of total Low Recall Positions (Out of 1,825,385) |
|---|---|---|
| LowCov (total) | 1,145,937 | 62.78% |
| - LowCov | 1,145,079 | 62.73% |
| - LowCov;Amb | 858 | 0.05% |
| Del (total) | 561,158 | 30.74% |
| - Del | 257,423 | 14.08% |
| - Del;LowCov | 302,826 | 16.55% |
| - Del;Amb | 719 | 0.04% |
| - Del;Amb;LowCov | 190 | 0.01% |
| PASS (Missed deletion)* | 113,848 | 6.24% |
| PASS (Genotype Disagreement with ground truth)** | 669 | 0.04% |
| Amb | 3,773 | 0.21% |

**Frequency of Pilon filter tags associated with positions with low empirical performance**. The distribution of variant filter annotations provided by the Pilon variant caller was evaluated across all positions which were not recalled correctly by Illumina WGS. ***Here the Illumina-based variant calling pipeline predicted a reference or alternate allele when instead a deletion was observed in the ground-truth assembly. ****Here there was disagreement on the base (reference or alternate) between the Illumina-based variant calling pipeline and the ground truth with no evidence for deletion.

#### Supplemental Table 5

| Pileup Mappability Parameters | % of PLC positions w/ P-map < 100% | % of non-PLC positions w/ P-map < 100% |
|---|---|---|
| K = 50 bp, E = 4 | 30.3% | 1.2% |
| K = 75 bp, E = 4 | 22.4% | 0.54% |
| K = 100 bp, E = 4 | 20.3% | 0.42% |
| K = 125 bp, E = 4 | 19.2% | 0.37% |
| K = 150 bp, E = 4 | 18.2% | 0.33% |

**The proportion of PLC and other regions with perfect P-map (across K = 50 bp to 150 bp).**

#### Supplemental Table 6

| Gene set | # genes | Median (+ IQR) mean-**P-Map (K=50,E=4)** score | # genes w/ mean- **P-Map (K=50,E=4)** < 1.0 |
|---|---|---|---|
| All other non-PLC genes | 3647 | 1.0000 (IQR: 1.0 - 1.0) | 231 (6.3%) |
| *pe/ppe* (PLC genes) | 168 | 0.9957 (IQR: 0.9676 - 1.0) | 110 (65.5%) |
| Mobile genetic elements (PLC) | 147 | 0.8779 (IQR: 0.1936 - 1.0) | 99 (67.3%) |
| Additional 69 repetitive genes* (PLC) | 69 | 0.9459 (IQR: 0.6267 - 0.9905) | 69 (100%) |

**Statistics from distribution of gene-level mean P-Map (K=50,E=4) scores between PLC and non-PLC regions**

Supplemental Figure 4.

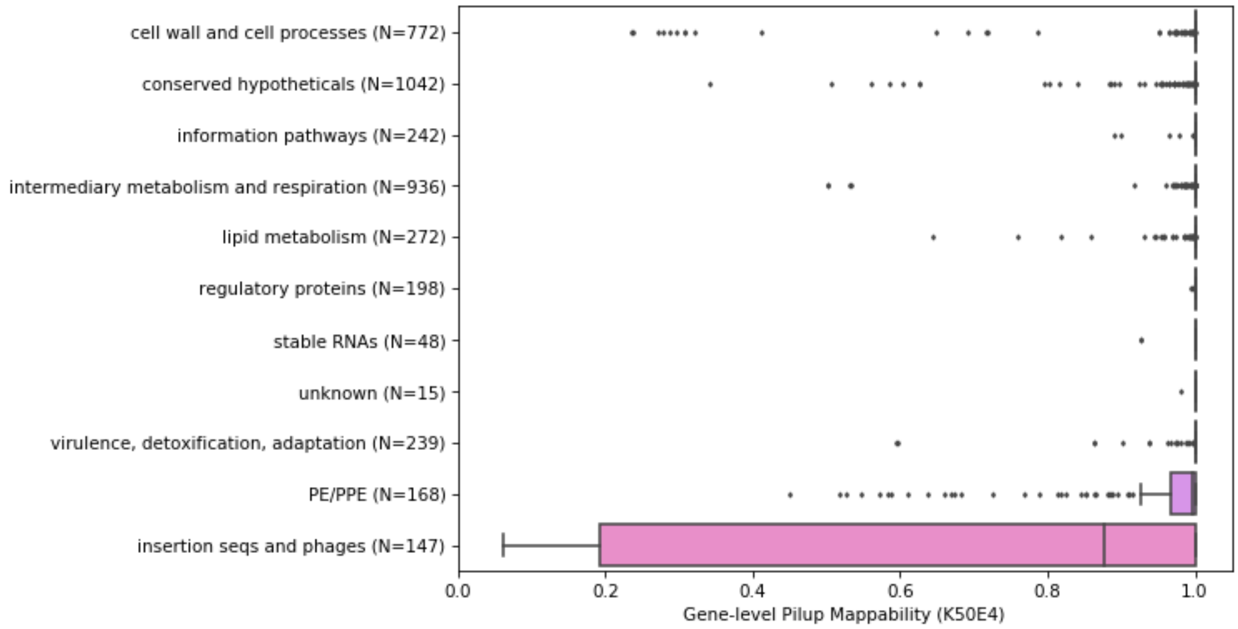


**Distribution of gene-level P-Map-K50E4 across all annotated functional gene categories in the Mtb genome**

#### Supplemental Table 7.

| Gene set | # genes | Median (+ IQR)<br>mean-**P-Map (K=50,E=4)** score | # genes w/ mean- **P-Map (K=50,E=4)** < 1.0 |
|---|---|---|---|
| cell wall and cell processes | 772 | 1.0000 (IQR: 1.0000 - 1.0000) | 63 (8.2% of total) |
| conserved hypotheticals | 1042 | 1.0000 (IQR: 1.0000 - 1.0000) | 68 (6.5% of total |
| information pathways | 242 | 1.0000 (IQR: 1.0000 - 1.0000) | 13 (5.4% of total |
| intermediary metabolism and respiration | 936 | 1.0000 (IQR: 1.0000 - 1.0000) | 73 (7.8% of total |
| lipid metabolism | 272 | 1.0000 (IQR: 1.0000 - 1.0000) | 46 (16.9% of total |
| regulatory proteins | 198 | 1.0000 (IQR: 1.0000 - 1.0000) | 10 (5.1% of total |
| Unknown | 15 | 1.0000 (IQR: 1.0000 - 1.0000) | 1 (6.7% of total |
| virulence, detoxification, adaptation | 239 | 1.0000 (IQR: 1.0000 - 1.0000) | 26 (10.9% of total |
| *pe/ppe* (PLC genes) | 168 | 0.9957 (IQR: 0.9676 - 1.0000) | 110 (65.5% of total |
| Mobile genetic elements (PLC) | 147 | 0.8779 (IQR: 0.1936 - 1.0000) | 99 (67.3% of total |

**Statistics from distribution of gene-level mean P-Map (K=50,E=4) across all annotated functional gene categories in the Mtb genome.**

Supplemental Figure 5

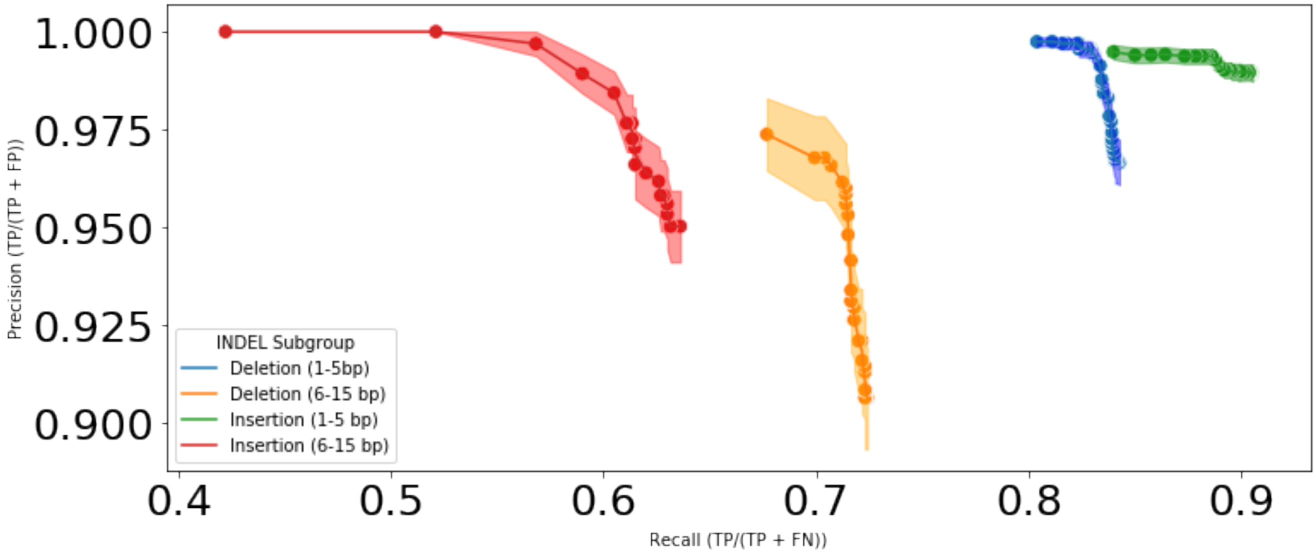


**INDEL detection performance stratified by length and type.** Splitting insertions and deletions into short (1-5bp) vs longer (6-15bp) variants, we found variant calling for the former to have performance comparable to SNSs. In contrast, detection of longer INDELs performed worse with much lower recall. Using a MQ filtering threshold of 40 the following performance was as follows: (1) short insertions (F1 = 0.94, precision=98.98%, recall=89.64%), (2) short deletions (F1 = 0.90, precision=99.43%, recall=83.05%). (3) longer insertions (F1 = 0.74, precision=96.61%, recall=61.49%), 4) longer deletions (F1 = 0.80, precision= 92.65%, recall = 71.79%). For all groups evaluated, the MQ thresholds evaluated ranged from 1-55. Complete benchmarking results can be found for each individual isolate in Additional File 11.

Supplemental Figure 6)

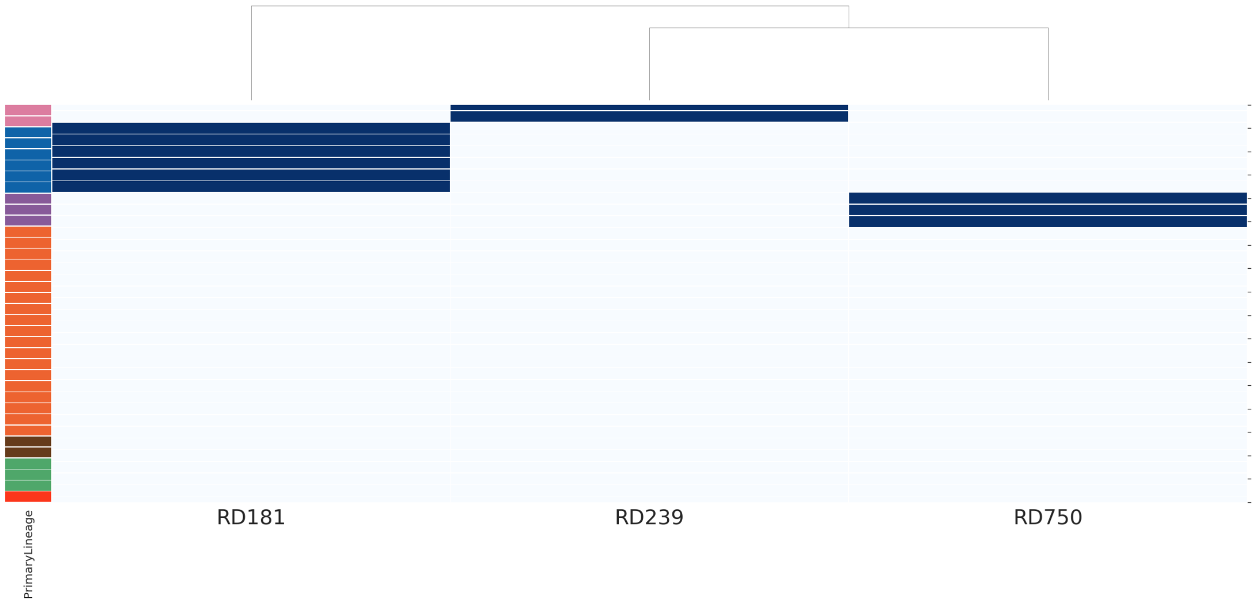


**Known regions of difference (RD) detected in lineages 1, 2, and 3.** The previously identified regions of difference (RD239, RD181, & RD750) associated with lineages 1, 2, and 3 were all identified in their expected lineage. None of the RDs were detected outside of their expected lineages. The vertical axis of the heatmap is colored by sample lineage (1-6 and then 8). Lineage 1 is colored pink, lineage 2 is colored blue, and lineage 3 is colored purple.

Supplemental Figure 7)

A

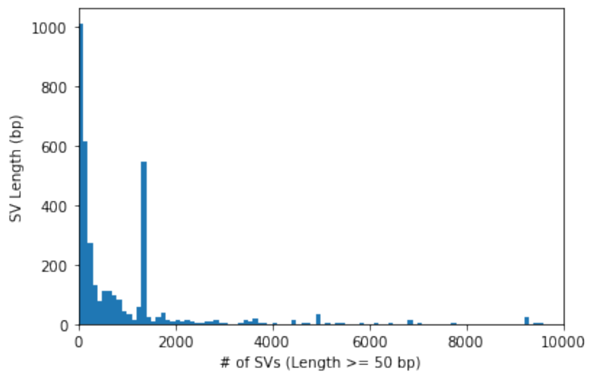


B

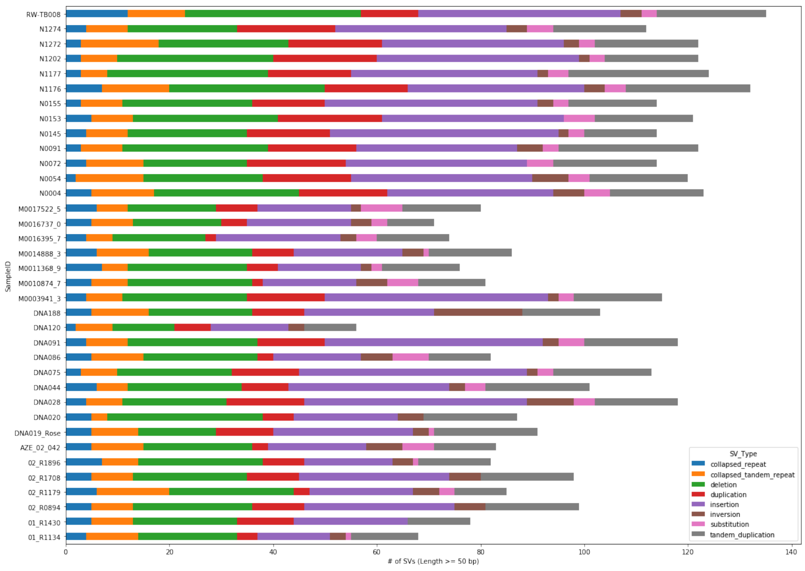


C

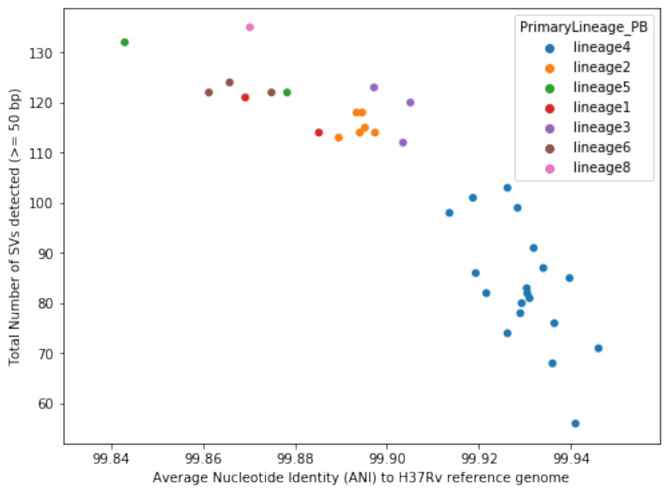


**Overview of structural variants detected across the 36 isolates.** A total of 3622 total SVs, with length ≥ 50 bp, were detected across all 36 isolates evaluated**. a)** We evaluated the distribution of SV lengths and found that the distribution was skewed towards a higher frequency of shorter SVs. The one exception was a spike in frequency around 1300-1350 bp, representing the higher frequency of SVs related to IS6110 type transposes in the Mtb genome. **b)** We evaluated the frequency of different SV classifications across all isolates evaluated. **c)** We evaluated the relationship between genetic distance, as measured by Average Nucleotide Identity, to the H37Rv reference sequence, and the # of SVs detected. There was a strong negative correlation between Average Nucleotide Identity to the reference, and the number of SVs detected relative to the reference (Spearman's $R = -0.899$, $p < 1.1e-13$). Each isolate is colored by the predicted MTBC lineage. The reference genome used, H37Rv, is a lineage 4 isolate.
